# An intact primordial interferon antiviral signalling network in amphioxus illuminates the basal chordate origin of vertebrate IFN immunity

**DOI:** 10.64898/2026.08.17.745176

**Authors:** Jingyuan Lin, Yilin Han, Huijuan Yang, Meng Yang, Guangdong Ji, Chen Sun, Zhenhui Liu

## Abstract

The evolutionary origin of the complete interferon (IFN) antiviral signalling cascade across chordates has long remained elusive, as fully functional IFN machinery had not been biochemically reconstituted in basal cephalochordates. Here we combine phylogenetics, AlphaFold3 structural prediction, multi-omic profiling and a full panel of *in vitro* and *in vivo* functional assays to characterize an intact primordial IFN network in amphioxus *Branchiostoma japonicum*, the extant sister group of all vertebrates. Amphioxus encodes a pair of undifferentiated BjTBK1/IKKε paralogs that form cytoplasmic heterodimers, whose conserved kinase-domain phosphorylation residues are mandatory for downstream IRF-driven IFN promoter activation. Despite minimal primary sequence homology with vertebrate IFNs, BjIFN1/2 possess compact α-helical core folds that hint at potential structural analogy to vertebrate mucosal type III IFN-λ. We identify two primitive class II cytokine receptors BjCRFB1 and BjCRFB2 that act as functional IFN sensors. Phylogenetic and tertiary structural validations confirm these undiversified ancestral receptor prototypes arose before vertebrate IFN receptor subfunctionalization, and their distinct temporal induction profiles upon viral challenge closely mirror the mucosal immune surveillance orchestrated by vertebrate IFN-λ–IFNLR signalling. AlphaFold3 ligand-receptor docking identifies BjIFN2–BjCRFB2 as the optimal binding pair among all tested combinations, yet all modelled complexes yield low interface scores indicative of weak ancestral intermolecular interactions. Downstream signal transduction relies on two STAT paralogs, BjSTATa and BjSTATb, which jointly mediate transcriptional activation of core antiviral effector genes. The conserved GTPase BjMx serves as a central IFN-stimulated effector to suppress viral replication and maintain tissue immune homeostasis. Collectively, our structural and functional evidence demonstrates that the full hierarchical IFN signalling apparatus was fully assembled in basal cephalochordates prior to vertebrate radiation. The amphioxus IFN cascade preserves core ancestral molecular traits including structure-dependent ligand conservation and primitive low-affinity ligand-receptor interactions. This study resolves a long-standing evolutionary gap and provides definitive functional evidence demonstrating that IFN-mediated innate immunity constitutes an ancestral chordate trait, rather than a vertebrate-specific evolutionary innovation. It also puts forward the hypothesis that mucosal surveillance may represent the ancestral mode of chordate IFN defence.

## Introduction

Innate interferon (IFN) signalling represents the primary and most diversified antiviral defence system in vertebrates. This multilayered, hierarchically regulated cascade orchestrates sequential immune events, including viral pattern recognition, upstream kinase activation, IFN ligand transcription and secretion, cell-surface receptor sensing, intracellular JAK-STAT signal transduction, and the final induction of interferon-stimulated genes (ISGs), thereby restricting viral propagation and restoring host immune homeostasis^1–3^. Cumulative mechanistic studies in mammalian and teleost models have extensively characterized the core composition and operational principles of vertebrate IFN signalling machinery^3–5^. Nevertheless, key evolutionary ambiguities remain unresolved: the phylogenetic origin of the complete IFN signalling toolkit during metazoan evolution, together with the ancestral structural and functional foundations that drove the emergence of sophisticated, subtype-specialized IFN networks in modern vertebrates, remain poorly defined.

Comparative genomic evidence has further revealed a distinct evolutionary boundary between protostome invertebrates and jawed vertebrates. Protostome lineages such as molluscs and arthropods encode partial JAK-STAT signalling components but lack canonical IFN ligands and specific IFN receptors, which precludes authentic IFN-dependent antiviral immunity in invertebrates^1^. By contrast, cartilaginous and bony fish harbour expanded IFN subtype repertoires and diversified receptor complexes; however, these evolutionarily derived vertebrate groups cannot recapitulate the undifferentiated ancestral state of chordate IFN immunity^5–7^. Most importantly, the full IFN signalling cascade in basal chordates has not yet been biochemically reconstituted or functionally verified, leaving the key evolutionary transition linking invertebrate innate immunity to canonical vertebrate IFN responses entirely uncharacterized.

As the sister group of all vertebrates, cephalochordate amphioxus retains conserved plesiomorphic genomic and anatomical features of early chordates, serving as an indispensable model for reconstructing the evolutionary origin of vertebrate innate immune systems^8–10^. Previous transcriptomic studies have predicted amphioxus IFN-like sequences and identified partial antiviral signalling molecules, including TLRs, RLRs and IRFs^10–16^. Despite these preliminary molecular annotations, current evidence is insufficient to support robust evolutionary inference. The antiviral activity of predicted amphioxus IFN proteins lacks rigorous *in vitro* and *in vivo* validation, and bona fide IFN receptors remain unidentified, leaving the ligand-sensing core of the chordate IFN cascade unconfirmed. In addition, a complete signalling axis that continuously bridges upstream viral detection and downstream ISG-mediated effector responses has not been experimentally reconstructed in amphioxus. These persistent gaps obscure the structural and functional integrity of basal chordate IFN networks, mask the ancestral prototypes of chordate IFN ligands and receptors, and limit our understanding of how primitive signalling molecules coordinated multi-step antiviral transduction before the gene duplication and subfunctionalization events that diversified vertebrate IFN systems.

To address these critical evolutionary immunological uncertainties, we systematically characterized core IFN cascade components in Branchiostoma japonicum through multi-omics profiling, structural bioinformatics analysis, and comprehensive functional immunological assays. We elucidate the full hierarchical regulatory framework of the primordial amphioxus IFN pathway, define conserved ancestral signatures and lineage-specific molecular features, and establish a stepwise evolutionary model of chordate IFN immunity. Our findings revise the prevailing view of vertebrate-specific *de novo* origin of complete IFN signalling, demonstrating instead that the full IFN signalling toolkit was already fully established in the most basal extant chordates.

## Results

### An undifferentiated TBK1/IKKε–IRF ancestral axis mediates virus-triggered IFN transcription

TBK1 and IKKε are non-canonical IKK kinases that serve as master upstream activators of vertebrate IFN production^17,18^, yet their undiversified ancestral configuration in basal chordates remains poorly characterized. We identified two paralogous TBK1/IKKε homologs in *B. japonicum*, designated BjTBK1/IKKε-a and BjTBK1/IKKε-b. Tertiary structure superimposition revealed high folding similarity between these two amphioxus kinases (Fig. 1A), which share comparable sequence homology to human and zebrafish TBK1 and IKKε (Supplementary Fig. S01). Maximum-likelihood phylogenetic reconstruction positioned BjTBK1/IKKε-a/b within the basal chordate clade, prior to the evolutionary separation of vertebrate TBK1 and IKKε subfamilies (Fig. 1B), supporting their identity as the undifferentiated ancestral precursors of vertebrate TBK/IKK kinases.

**Fig. 1.**
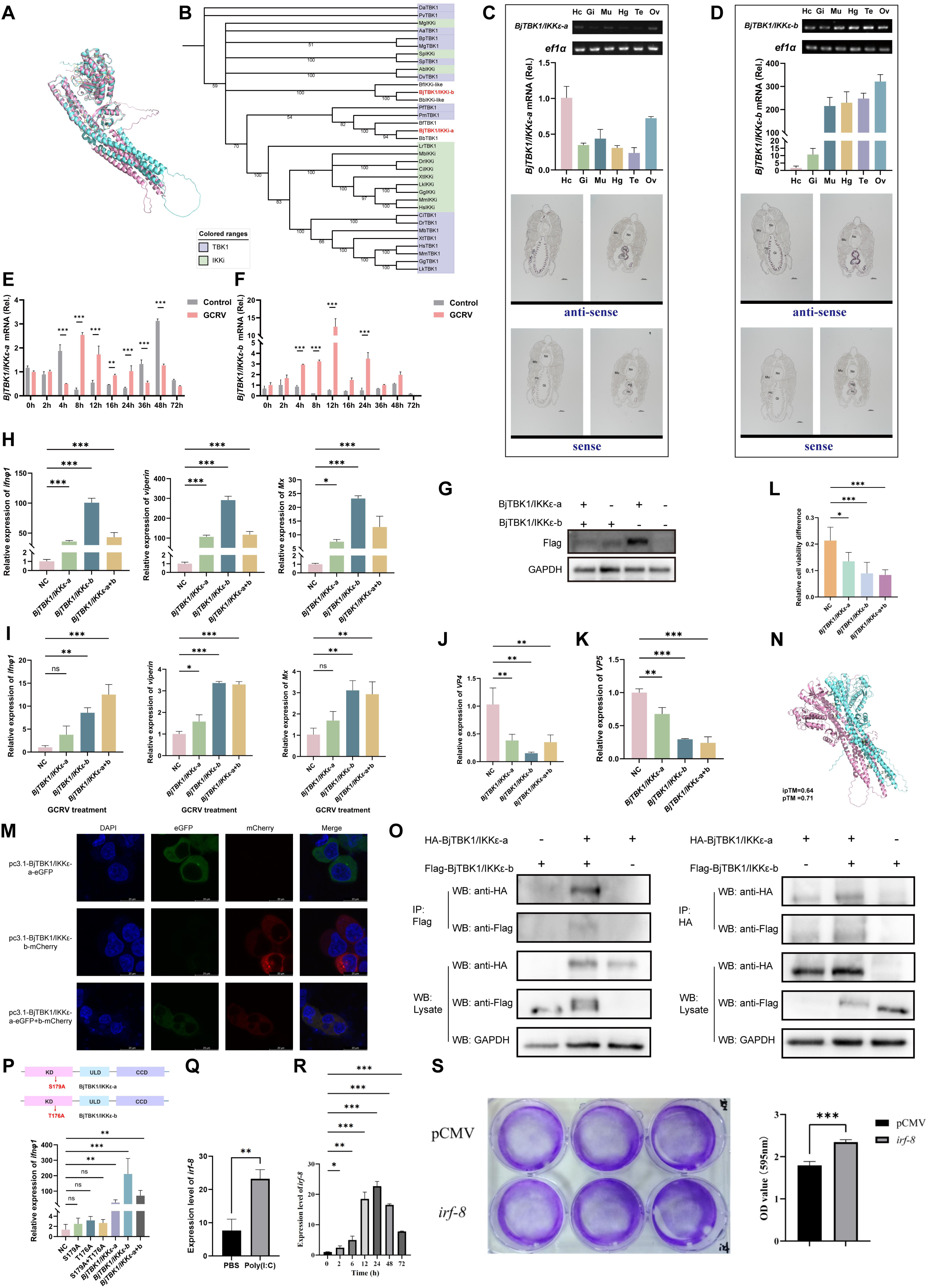
An ancestral TBK1/IKKε–IRF cascade mediates virus-induced IFN transcription in amphioxus. **(A)** Homology-based structural modelling of BjTBK1/IKKε-a and BjTBK1/IKKε-b from *B. japonicum*. **(B)** Maximum-likelihood phylogenetic reconstruction of TBK1/IKKε family proteins. BjTBK1/IKKε-a and BjTBK1/IKKε-b cluster within the basal chordate clade, preceding the evolutionary divergence of vertebrate TBK1 and IKKε subfamilies. **(C,D)** Tissue-wide expression profiles of BjTBK1/IKKε-a and BjTBK1/IKKε-b in adult amphioxus. Top panels: semi-quantitative RT-PCR; middle panels:qRT-PCR; bottom panels: *in situ* hybridisation, showing antisense and sense probe controls. Hc, hepatic caecum; Gi, gill; Mu, muscle; Hg, hindgut; Te, testis; Ov, ovary. **(E,F)** Time-course qRT-PCR analysis of *BjTBK1/IKKε-a* and *BjTBK1/IKKε-b* upon GCRV challenge. Both paralogs are significantly transcriptionally upregulated across multiple time-points post-virus infection. **(G-I)** Functional overexpression assays in EPC cells. Ectopic expression of either BjTBK1/IKKε-a or BjTBK1/IKKε-b induces transcription of fish *ifnφ1*, *viperin* and *Mx*, with further elevated expression upon GCRV infection. Immunoblot in (G) confirms protein overexpression. (J,K) qRT-PCR quantification of GCRV *vp4* and *vp5* viral transcripts. Overexpression of BjTBK1/IKKε paralogs represses viral gene transcription following GCRV stimulation. (L) CCK-8 cell viability assay. EPC cells overexpressing BjTBK1/IKKε-a or BjTBK1/IKKε-b show improved survival against GCRV-triggered cytopathic injury relative to empty vector control cells. **(M)** Subcellular localisation analysis. BjTBK1/IKKε-a and BjTBK1/IKKε-b localize predominantly to the cytoplasm and display extensive co-localization, assessed by confocal immunofluorescence microscopy. **(N)** AlphaFold-predicted structural model for the heteromeric complex formed by BjTBK1/IKKε-a and BjTBK1/IKKε-b. **(O)** Reciprocal co-immunoprecipitation (Co-IP) verifying direct physical interaction between BjTBK1/IKKε-a and BjTBK1/IKKε-b in transfected EPC cells. Whole cell lysates are shown as input controls. **(P)** qRT-PCR measurement of *ifnφ1* induction driven by wild-type and phosphorylation-deficient BjTBK1/IKKε mutants. Mutation of conserved kinase-domain phosphorylation residues (S179A for BjTBK1/IKKε-a; T176A for BjTBK1/IKKε-b) abolishes the capacity to activate *ifnφ1* transcription. **(Q)** qRT-PCR analysis of *Bjirf-8* in amphioxus intestinal tissue following Poly(I:C) challenge. Amphioxus were intraperitoneally injected with Poly(I:C) (0.02 μg/μL, dissolved in PBS), while control animals received an equal volume of PBS. Intestinal tissues were harvested 24 h post-injection for transcript quantification. **(R)** Time-course qRT-PCR detection of *Bjirf-8* after GCRV intraperitoneal injection. Tissue samples were collected at 0 h, 2 h, 6 h, 12 h, 24 h, 48 h and 72 h post-challenge. *ef1α* was used as the internal reference gene. **(S)** Crystal violet staining-based cellular reporter assay assessing *Bjirf-8*-dependent cellular responses; absorbance was measured at OD_595_ nm. All quantitative data represent mean ± SEM from three independent biological replicates. Statistical significance was determined by one-way ANOVA or Student’s *t*-test; *\*P* < 0.05, *\*\*P* < 0.01, *\*\*\*P* < 0.001.

Tissue-wide qPCR profiling uncovered overlapping expression of both paralogs in primary immune tissues, including the gill, hepatic caecum and hindgut, alongside biased enrichment in reproductive and muscle tissues (Fig. 1C, D). Time-course qPCR assays detected robust transcriptional upregulation of both genes at multiple time points following GCRV infection (Fig. 1E, F). Heterologous overexpression in EPC cells further validated their antiviral activity: individual overexpression of either BjTBK1/IKKε paralog strongly induced fish *ifnφ1* and its downstream interferon stimulated genes (*viperin*, *Mx*) (Fig. 1G-I), repressed transcript levels of the GCRV structural genes *vp4* and *vp5* (Fig. 1J, K), and alleviated virus-triggered cytopathic effects (Fig. 1L).

Confocal immunofluorescence demonstrated that both kinases localize exclusively to the cytoplasm and exhibit full co-localization (Fig. 1M). Reciprocal co-immunoprecipitation (Co-IP) experiments confirmed direct physical assembly of heteromeric kinase complexes, consistent with interaction interfaces predicted by AlphaFold modelling (Fig. 1N, O). Multiple sequence alignment identified deeply conserved serine/threonine phosphorylation sites within the kinase domain, corresponding to Ser^179^ in BjTBK1/IKKε-a and Thr^176^ in BjTBK1/IKKε-b (Supplementary Fig. S02). Site-directed mutagenesis at these residues abolished the capacity of the kinases to drive *ifnφ1* transcription in EPC cells (Fig. 1P), establishing phosphorylation-dependent activation as an evolutionarily conserved regulatory switch.

Genome-wide homology searches retrieved multiple amphioxus proteins harbouring canonical IRF3-type domains, which group phylogenetically with vertebrate IRF3/7 homologues (Supplementary Fig. S03, S04). Among these IRF family members, BjIRF8 exhibited robust transcriptional induction in intestinal tissue upon poly(I:C) or GCRV challenge (Fig. 1Q, R). Gain-of-function experiments in EPC cells further confirmed that BjIRF8 confers potent cytoprotection against viral insult (Fig. 1S).

Earlier studies have described the TBK1-IRF3/7 phosphorylation-dependent regulatory cascade in amphioxus^11^. By integrating these prior observations with our functional characterizations, we establish a complete hierarchical model for the amphioxus primordial antiviral signalling circuit: upon viral stimulation, cytoplasmic BjTBK1/IKKε heterodimers are phosphorylated; activated kinase complexes physically interact with IRF3/7- like proteins and trigger their serine phosphorylation; phosphorylated IRF proteins translocate to the nucleus and cooperatively initiate transcription of primitive chordate IFN ligands.

### Identification of Amphioxus IFN-like ligands

Extreme interspecies sequence divergence has long complicated the identification of deeply diverged IFN homologs in invertebrate chordates, because conventional sequence-based homology searches lose sensitivity when primary-sequence conservation is low. We therefore hypothesized that evolutionarily conserved structural features of IFNs may be retained more strongly than primary-sequence similarity. To identify candidate amphioxus IFN-like proteins, we established a hierarchical multimodal screening workflow integrating transcriptome-derived protein filtering, secretory-protein prediction, domain annotation, AlphaFold-based structural prediction, Foldseek-based structural similarity searches, and subsequent sequence and phylogenetic evaluation (Fig. 2A). A non-redundant predicted protein dataset was generated from the transcriptomic data of *B. japonicum*, and the integrity of the predicted open reading frames (ORFs) was systematically evaluated. To minimize the loss of potential candidates through overly restrictive size filtering, proteins ≤300 amino acids in length were preferentially retained for subsequent screening. SignalP and DeepTMHMM were then used to predict N-terminal signal peptides and transmembrane topology, respectively, and proteins containing an intact signal peptide but lacking a typical transmembrane domain were prioritized as putative secreted cytokine-like proteins. InterProScan and Pfam analyses were subsequently used for protein-family and conserved-domain annotation, allowing sequences assigned to well-defined enzymes, membrane receptors, or other non-cytokine protein classes to be excluded. The remaining candidates were subjected to three-dimensional structural prediction, and proteins displaying compact, α-helix-rich cytokine-like folds were retained for further analysis. Foldseek was then used for structure-based similarity searches against known IFN proteins. Candidates were retained when at least one structural hit met all predefined thresholds: E-value ≤1, TM-score ≥0.4, query coverage ≥0.7, and target coverage ≥0.5. Sequences passing this structural screen were further examined using NCBI BLASTP as a complementary sequence-based assessment.

**Fig. 2.**
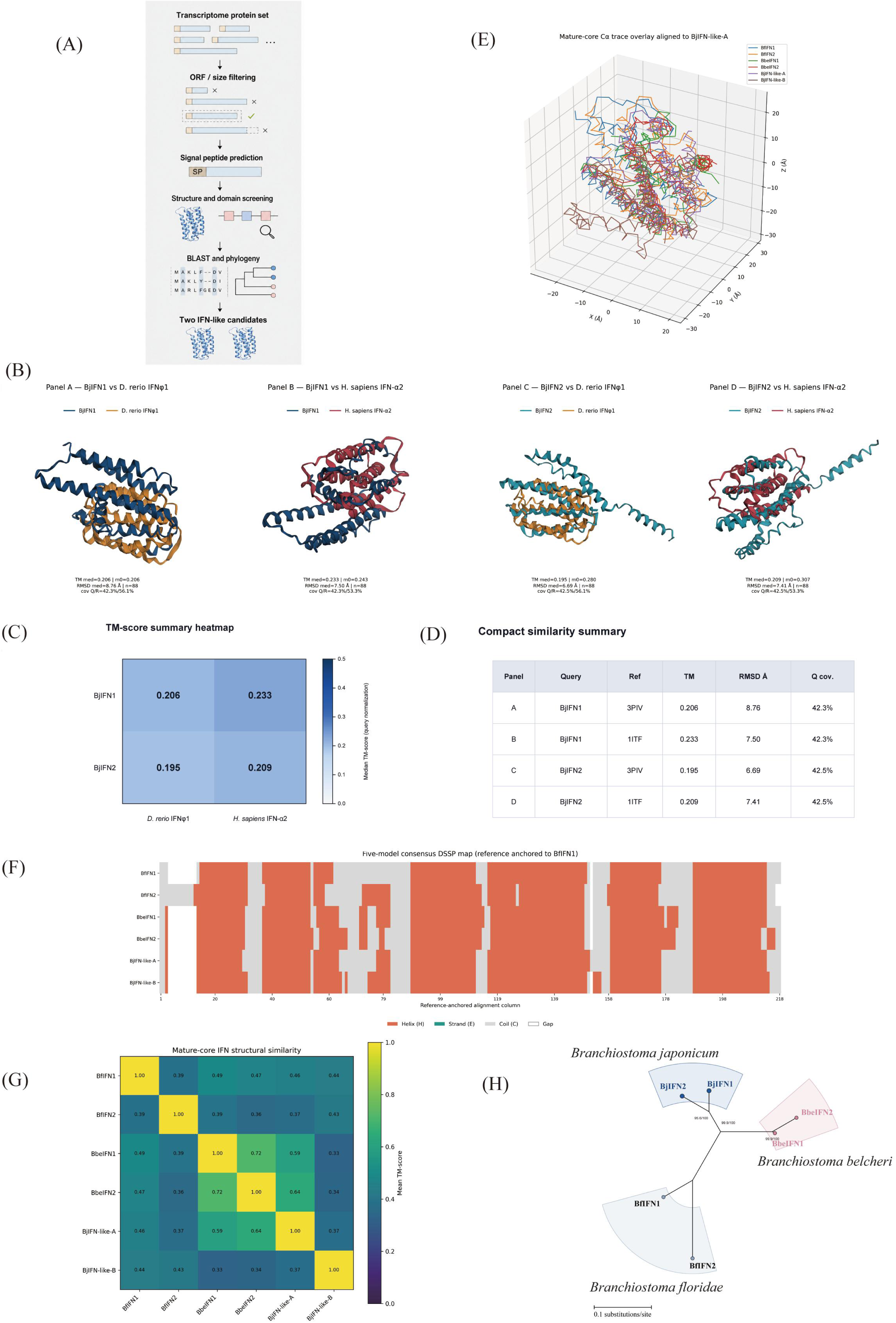
Identification and comparative structural analysis of amphioxus IFN-like proteins. **(A)** Schematic workflow for hierarchical screening of IFN-like candidates from the *B. japonicum* transcriptome, incorporating ORF and size filtering, signal-peptide prediction, structural and domain screening, and BLAST- and phylogeny-based evaluation. **(B)** Pairwise structural superimpositions of BjIFN1 and BjIFN2 with vertebrate interferons: BjIFN1 versus zebrafish IFNφ1 (PDB 3PIV), BjIFN1 versus human IFN-α2 (PDB 1ITF), BjIFN2 versus zebrafish IFNφ1, and BjIFN2 versus human IFN-α2. Corresponding TM-score, RMSD, and query-coverage statistics are indicated beneath each comparison. **(C)** TM-score heatmap summarizing the four BjIFN–vertebrate IFN structural comparisons shown in panel B. **(D)** Compact summary of TM-score, RMSD, and query-coverage values obtained from the pairwise structural alignments. The relatively low global TM-scores indicate limited overall structural similarity between amphioxus and vertebrate IFNs, consistent with substantial evolutionary divergence. **(E)** Superimposition of mature-core Cα traces from amphioxus IFN-like proteins after structural alignment to BjIFN1, illustrating similarities and differences in their overall core architectures. **(F)** DSSP-based secondary-structure map of mature amphioxus IFN-like proteins after reference-anchored alignment, showing the distribution of α-helical and other secondary-structure elements across aligned positions. **(G)** Pairwise TM-score heatmap of mature-core structural similarity among amphioxus IFN-like proteins. **(H)** Maximum-likelihood phylogenetic tree of IFN-like proteins from *B. japonicum*, *B. belcheri*, and *B. floridae*. Node support values are indicated at internal branches, and the scale bar represents 0.1 substitutions per site.

Two proteins, designated BjIFN1 (GenBank PZ746091) and BjIFN2 (GenBank PZ746092), satisfied the combined secretory, domain, and structural screening criteria. Pairwise structural superimposition of BjIFN1 and BjIFN2 with zebrafish IFNφ1 and human IFN-α2 yielded relatively low global TM-scores (0.195–0.233), indicating limited overall structural similarity to the vertebrate IFN reference structures (Fig. 2B–D). Nevertheless, both BjIFN candidates retained compact, α-helix-rich core architectures, while comparative analyses of amphioxus IFN-like proteins further revealed recurrent helical organization and measurable structural conservation within their mature-core regions (Fig. 2E-G). Furthermore, structural superimposition and DSSP secondary-structure profiling based on AlphaFold-predicted models revealed that BjIFN1 and BjIFN2 do not closely reproduce the long, relatively straight helical arrangement characteristic of vertebrate type I IFNs (Fig. 2B, F). Instead, BjIFN1 and BjIFN2 adopt a streamlined monomeric helical core architecture, which may suggest potential structural resemblance to vertebrate type III IFN-λ (Fig. 2G).

Vertebrate IFNs are broadly classified into four types (Types I–IV)^19^. However, the extensive sequence divergence among IFNs complicates subtype assignment based solely on primary-sequence similarity or phylogenetic inference. Comparative analysis of IFN-like proteins from three amphioxus species (*B. japonicum*, *B. floridae*, and *B. belcheri*) revealed relatively high within-genus full-length sequence identities (71.7–94.7%) and broadly similar secondary-structure compositions. α-Helices accounted for 68.85–77.60% of the mature-core regions, indicating substantial conservation of helical organization among amphioxus IFN-like proteins (Fig. 2F; Supplementary Table S1; Supplementary Figs. S05 and S06). Structural superimposition of the mature-core Cα traces, together with pairwise TM-score comparisons, further demonstrated conservation of a common helical-core architecture despite variation in peripheral regions (Fig. 2E, G). Maximum-likelihood phylogenetic analysis of IFN-like proteins from the three amphioxus species revealed species-associated clustering patterns consistent with lineage-specific duplication and diversification within cephalochordates (Fig. 2H).

Taken together, primitive chordate IFN ligands rely on conserved α-helical tertiary architecture rather than linear sequence similarity to maintain biological function, and BjIFN1/2 represent the structural evolutionary progenitors of all vertebrate interferons.

### BjCRFB1 and BjCRFB2 exhibit structural and antiviral functional features of primitive IFN receptors

In vertebrates, IFN signalling is initiated by dedicated cell-surface class II cytokine receptors that mediate ligand recognition and downstream signal transduction^20^. Prior to this study, no functionally characterized IFN-receptor-like molecule had been established in amphioxus, leaving an important gap in understanding the evolutionary origin of the vertebrate IFN receptor system. In teleosts, type I IFN receptor chains belong to the cytokine receptor family B (CRFB)^21,22^. Zebrafish CRFB1 and CRFB2 function as counterparts of mammalian IFNAR2, whereas CRFB5 represents the shared receptor chain corresponding functionally to mammalian IFNAR1^23^. Searches using zebrafish CRFB5 did not recover a convincing amphioxus homologue. In contrast, searches using zebrafish CRFB1 and CRFB2 identified two CRFB-related genes in the *B. japonicum* genome, which we designated BjCRFB1 and BjCRFB2. Phylogenetic reconstruction based on their extracellular-domain sequences placed both proteins within the broader class II cytokine receptor lineage (Supplementary Fig. S07), supporting an evolutionary relationship between amphioxus CRFBs and vertebrate cytokine receptor systems. Notably, BjCRFB1 and BjCRFB2 occupied distinct positions within this lineage, suggesting that structural diversification of the two amphioxus receptors may have occurred early in chordate evolution.

Domain-architecture analysis further revealed canonical class II cytokine receptor features in both proteins, including an N-terminal signal peptide, extracellular fibronectin type III (FN3) domains and a single transmembrane segment. BjCRFB1 contains four tandem FN3 domains, whereas BjCRFB2 contains three. Comparison with human IFNAR1 and IFNAR2 demonstrated broadly similar receptor domain architectures (Fig. 3A), with additional vertebrate class II cytokine receptor comparisons shown in Supplementary Fig. S08. To assess tertiary-structure conservation, BjCRFB1 and BjCRFB2 were structurally superimposed with representative human and zebrafish class II cytokine receptors (Fig. 3B). BjCRFB1 showed fold-level similarity to human IL10RA and zebrafish CRFB5, with TM-scores of 0.40321 and 0.39735, respectively, whereas BjCRFB2 showed higher structural similarity to human IFNGR2 and zebrafish CRFB5, with TM-scores of 0.53364 and 0.57097, respectively. Together with the broader structural comparisons (Supplementary Figs. S09 and S10; Supplementary Tables S2 and S3), these results indicate that BjCRFB1 and BjCRFB2 retain recognizable class II cytokine receptor folds while exhibiting distinct structural affinities toward different vertebrate receptor subfamilies.

**Fig. 3:**
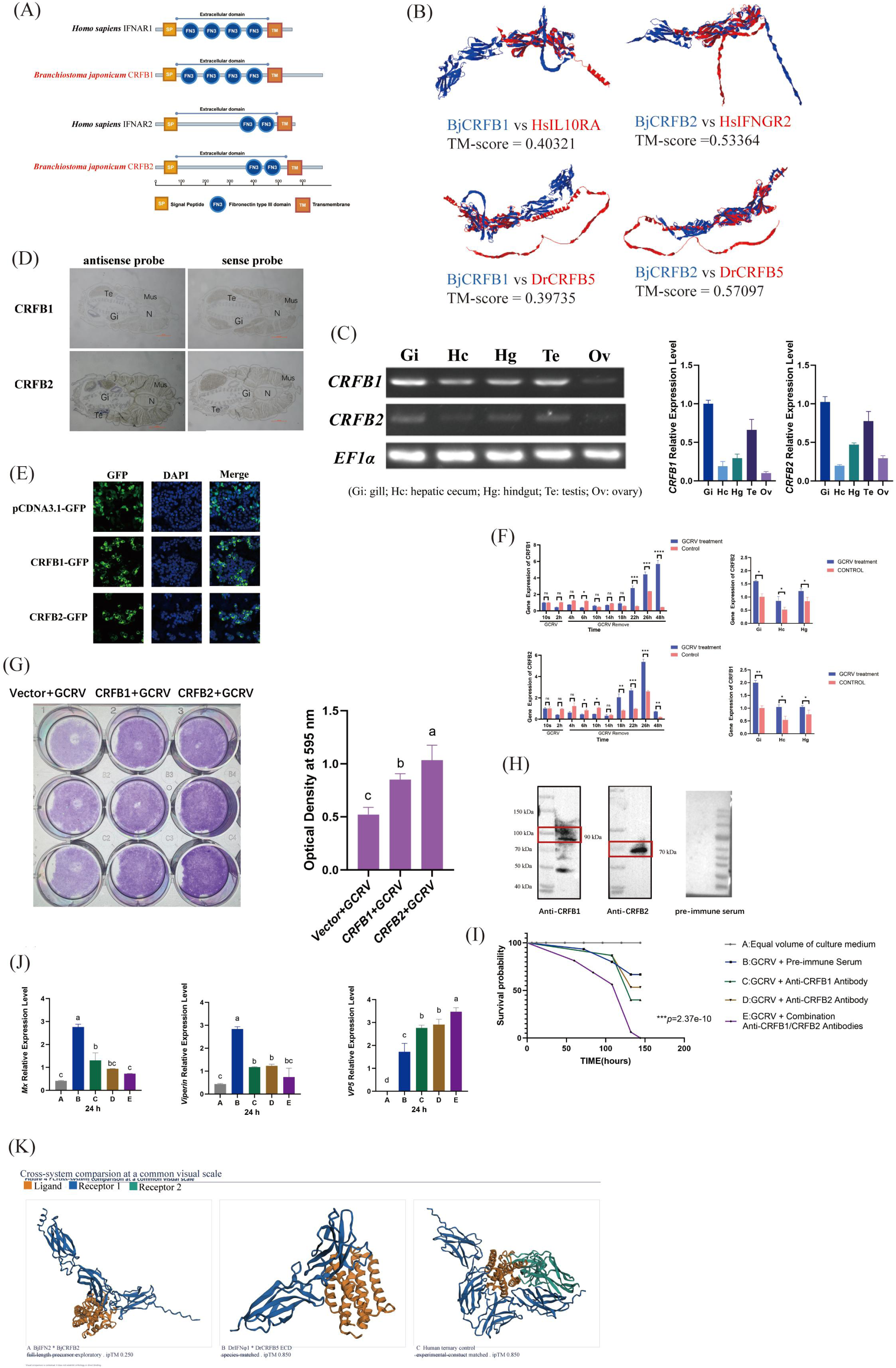
BjCRFB1 and BjCRFB2 function as primitive IFN receptors governing amphioxus antiviral signalling. **(A)** Domain architectures of *B. japonicum* BjCRFB1 and BjCRFB2 compared with human IFNAR1 and IFNAR2. Signal peptides (SP), extracellular fibronectin type III (FN3) domains, and transmembrane regions (TM) are indicated. **(B)** Three-dimensional structural superimposition of BjCRFB1 and BjCRFB2 with representative vertebrate class II cytokine receptors. BjCRFB1 was compared with human IL10RA and zebrafish CRFB5, whereas BjCRFB2 was compared with human IFNGR2 and zebrafish CRFB5. Corresponding TM-scores are shown beneath each comparison. BjCRFB proteins are shown in blue and vertebrate reference receptors in red, illustrating varying degrees of global structural similarity. **(C)** Tissue expression profiles of *BjCRFB1* and *BjCRFB2*. Left, semi-quantitative RT-PCR analysis with *EF1α* as the internal control; right, qRT-PCR quantification of relative transcript abundance. Gi, gill; Hc, hepatic caecum; Hg, hindgut; Te, testis; Ov, ovary. **(D)** *In situ* hybridization analysis of *BjCRFB1* and *BjCRFB2* expression. Antisense probes were used to detect endogenous transcripts, with corresponding sense probes serving as negative controls. **(E)** Subcellular localization of BjCRFB1 and BjCRFB2. Cells expressing pCDNA3.1-GFP, BjCRFB1-GFP, or BjCRFB2-GFP were examined by fluorescence microscopy. GFP signals indicate the localization of the fusion proteins, DAPI marks nuclei, and merged images show signal distribution relative to the nucleus. **(F)** Transcriptional responses of *BjCRFB1* and *BjCRFB2* following grass carp reovirus (GCRV) infection. Left, time-course qRT-PCR analysis after infection; right, tissue-specific expression changes after GCRV challenge. **(G)** Effects of BjCRFB1 or BjCRFB2 overexpression on GCRV-induced cytopathic injury in EPC cells. Left, crystal-violet staining of GCRV-infected cells; right, quantification of retained crystal violet by optical density at 595 nm (OD595). **(H)** Immunoblot validation of polyclonal antibodies against BjCRFB1 and BjCRFB2. Antibody reactivity against the corresponding target proteins was assessed, with pre-immune serum included as a negative control. **(I)** Kaplan–Meier survival analysis of GCRV-infected amphioxus following antibody-mediated receptor blockade. Survival was compared among culture-medium control, GCRV plus pre-immune serum, GCRV plus anti-BjCRFB1 antibody, GCRV plus anti-BjCRFB2 antibody, and GCRV plus combined anti-BjCRFB1/anti-BjCRFB2 treatment groups. **(J)** Effects of BjCRFB1/2 blockade on antiviral gene expression and viral burden in amphioxus gills. Relative transcript levels of the antiviral effector genes *Mx* and *Viperin*, together with the GCRV structural gene *VP5*, were quantified by qRT-PCR. **(K)** Cross-species structural comparison of amphioxus, zebrafish, and human IFN–receptor complexes displayed at a common visual scale. Left, exploratory AlphaFold3 prediction of the BjIFN2–BjCRFB2 binary complex; middle, zebrafish DrIFNφ1–DrCRFB5 extracellular-domain complex; right, the human IFN receptor ternary system used as a positive structural reference. Ligands are shown in orange, the primary receptor chain in blue, and the secondary receptor chain in green. ipTM values are indicated for predicted complexes.

Tissue-expression profiling showed that both BjCRFB1 and BjCRFB2 were expressed across multiple amphioxus tissues, with comparatively high expression in immune-associated tissues including the gill and hindgut (Fig. 3C). *In situ* hybridization further supported their tissue distribution patterns (Fig. 3D). Fluorescence imaging of BjCRFB1-GFP and BjCRFB2-GFP fusion proteins revealed prominent localization at the plasma membrane, consistent with their predicted roles as cell-surface receptors (Fig. 3E). We previously demonstrated that grass carp reovirus (GCRV) can cross species barriers to infect amphioxus^24^. Following GCRV challenge, transcription of both genes was induced, but the two receptors displayed distinct temporal response profiles (Fig. 3F). BjCRFB2 responded earlier following infection, whereas BjCRFB1 exhibited a more delayed and sustained induction. Tissue-specific analysis at 24 h post-infection further showed that the strongest transcriptional responses of both receptors occurred in the gill.

We next examined whether BjCRFB1 and BjCRFB2 contribute functionally to antiviral defence. Exploratory structural modelling had previously suggested potential compatibility of their extracellular domains with fish IFNφ-related ligands and of their intracellular regions with JAK–STAT-associated signalling proteins, although these predicted interfaces were of relatively low confidence (Supplementary Fig. S11). In EPC cells, overexpression of either BjCRFB1 or BjCRFB2 reduced GCRV-induced cytopathic loss, as assessed by crystal-violet staining, with BjCRFB2 producing the stronger protective phenotype (Fig. 3G). To evaluate receptor function *in vivo*, polyclonal antibodies targeting the extracellular regions of BjCRFB1 and BjCRFB2 were generated and validated by immunoblotting (Fig. 3H). Antibody-mediated blockade of either receptor reduced the survival of GCRV-infected amphioxus, whereas simultaneous blockade of both receptors produced the most pronounced survival defect (Fig. 3I). Consistent with these phenotypic changes, receptor blockade reduced transcription of the antiviral effector genes *Mx* and *Viperin* while increasing the abundance of the viral structural gene *VP5* in gill tissues, with the combined blockade producing the strongest effect (Fig. 3J). These complementary gain- and loss-of-function experiments demonstrate that BjCRFB1 and BjCRFB2 both contribute to antiviral defence and possess partially overlapping antiviral functions.

To place the predicted amphioxus ligand–receptor interaction in an evolutionary structural context, we compared the BjIFN– BjCRFB system with representative zebrafish and human IFN receptor complexes at a common visual scale (Fig. 3K). Among the four exploratory BjIFN–BjCRFB combinations examined by AlphaFold3, the BjIFN2–BjCRFB2 model displayed the relatively most favourable predicted interface geometry (Supplementary Fig. S14) and was therefore selected for cross-system visualization. This binary amphioxus model was compared with the zebrafish DrIFNφ1–DrCRFB5 extracellular-domain complex and a human IFN–IFNAR ternary receptor system used as a vertebrate structural reference. The BjIFN2–BjCRFB2 model exhibited a plausible ligand–receptor arrangement but relatively low interface confidence, consistent with a weakly constrained or evolutionarily primitive interaction geometry. These modelling results support structural compatibility between amphioxus IFN-like ligands and BjCRFB receptors.

To further characterize the downstream regulatory programmes associated with BjCRFB1 and BjCRFB2, we performed transcriptomic profiling of amphioxus gills following antibody-mediated blockade of either receptor during GCRV infection. KEGG enrichment analysis revealed distinct but overlapping immune-related pathways following BjCRFB1 or BjCRFB2 inhibition. BjCRFB1 blockade preferentially affected pathways associated with necroptosis, neutrophil extracellular trap formation and related cellular programmes, whereas BjCRFB2 blockade prominently affected NOD-like receptor signalling, MAPK signalling and apoptosis-related pathways (Supplementary Fig. S15). qRT-PCR validation further showed that blockade of either receptor altered expression of shared JAK–STAT-associated components, including Jak2, Stata and Statb, while also producing receptor-specific effects on additional immune and cell-death-related genes (Supplementary Fig. S16). These results indicate that BjCRFB1 and BjCRFB2 share a common antiviral signalling core while regulating partially divergent downstream transcriptional programmes.

Collectively, the phylogenetic, domain-architecture, structural, expression and functional data support BjCRFB1 and BjCRFB2 as antiviral class II cytokine receptor-like components with features consistent with a primitive IFN receptor system in amphioxus. Their plasma-membrane localization, virus-responsive expression, antiviral gain- and loss-of-function phenotypes, and association with JAK–STAT-related transcriptional regulation establish a functional link between these receptors and amphioxus antiviral immunity. Together with the predicted structural compatibility between BjIFN-like ligands and BjCRFB receptors, these findings provide evidence for an early chordate stage in the evolution of vertebrate IFN receptor signalling.

### A flexible ancestral JAK–STAT cascade transduces IFN downstream antiviral signals

Vertebrate STAT paralogs undergo strict subfunctionalization for specialized IFN signalling roles^25,,26^, yet their undifferentiated ancestral state in basal chordates remained uncharacterized. Here, we identified two STAT paralogs in amphioxus *B. japonicum*, designated BjSTATa and BjSTATb. Phylogenetic analysis positioned BjSTATa basal to the vertebrate STAT1–STAT4 subfamily, and BjSTATb at the root of the vertebrate STAT5 clade (Supplementary Fig. S17), demonstrating vertebrate STAT subtype diversification occurred post-chordate divergence.

Tissue expression profiling showed that *BjSTATa* and *BjSTATb* are ubiquitously expressed across amphioxus tissues, with significant enrichment in immune tissues including gill, hepatic caecum and hindgut (Fig. 4A, B). Time-series qPCR assays demonstrated that both genes were significantly upregulated from 4 h post GCRV infection and maintained sustained elevated expression within 72 h (Fig. 4C, D). Subcellular imaging revealed divergent trafficking: both paralogs localized to the cytoplasm at resting state; viral stimulation triggered robust nuclear translocation exclusively for BjSTATb, marking it the primary transcriptional effector (Fig. 4E). We further evaluated the antiviral capacity of BjSTATa and BjSTATb via cytopathic effect (CPE) observation, crystal violet staining and viral gene quantification. Overexpression of BjSTATa or BjSTATb markedly alleviated GCRV-induced cell injury, and co-overexpression of the two paralogs produced enhanced protective effects (Fig. 4F). Consistent with the phenotypic rescue, the transcript levels of viral structural genes *vp4* and *vp5* were significantly decreased upon overexpression of BjSTATa and/or BjSTATb (Fig. 4G).

**Fig. 4:**
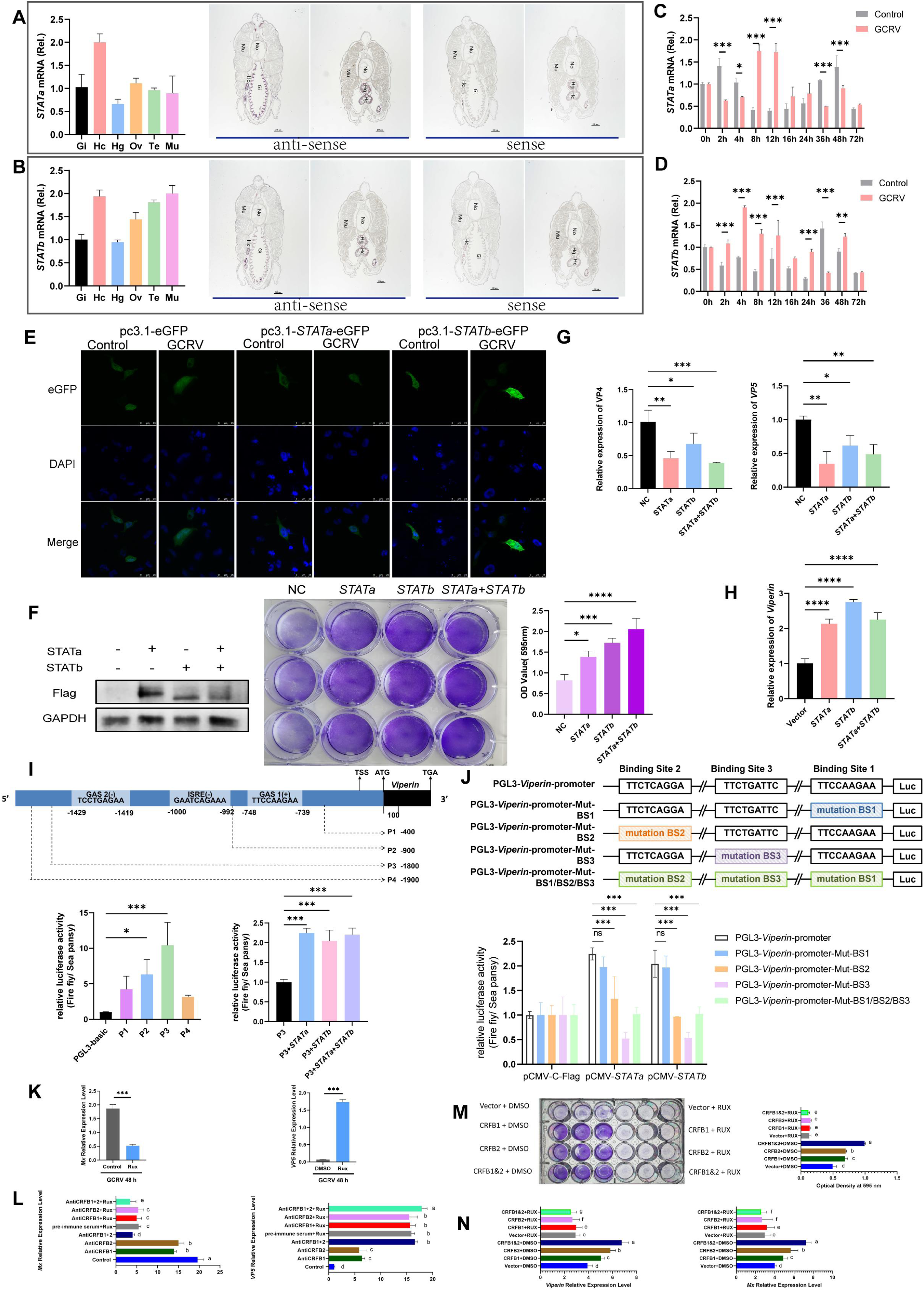
A primitive JAK-STAT pathway transduces IFN-dependent antiviral signals in amphioxus. **(A, B)** Tissue-distribution profiles of *BjSTATa* and *BjSTATb* mRNA across amphioxus tissues, determined by qRT-PCR (left) and *in situ* hybridization (right). Gill, hepatic caecum and hindgut show enriched transcript abundance for both paralogs. **(C, D)** Time course mRNA expression of *BjSTATa* and *BjSTATb* following GCRV viral challenge. **(E)** Subcellular localization of BjSTATa-eGFP and BjSTATb-eGFP under resting and virus-stimulated conditions. (F, G) Antiviral phenotypes upon BjSTATa/BjSTATb overexpression in EPC cells. **(F)** Crystal-violet staining and quantitative cytoprotection assays following GCRV infection. Individual overexpression of BjSTATa or BjSTATb alleviates virus-induced cytopathic effect (CPE). **(G)** qRT-PCR quantification of viral *vp4* and *vp5* transcript levels. BjSTAT overexpression reduces viral-gene abundance. **(H)** qRT-PCR measurement of *viperin* mRNA induction upon BjSTATa or BjSTATb overexpression at 4 h post-GCRV infection. **(I)** Dual-luciferase reporter assays testing serial truncated fragments of the *viperin* promoter. **(J)** Site-mutagenesis analysis of three predicted STAT-binding sites (BS1-BS3) within the P3 promoter region. **(K, L)** *In vivo* pharmacological blockade assays in amphioxus. **(K)** Effects of JAK inhibitor ruxolitinib (RUX) treatment: RUX suppresses antiviral *Mx* expression and increases GCRV *vp5* viral loads in gill tissue. **(L)** Combined neutralization of BjCRFB1/2 plus RUX co-treatment yields additive impairment of antiviral responses, further repressing Mx and elevating viral loads relative to single-agent treatments. **(M, N)** *In vitro* EPC cell gain-of-function validation of pathway hierarchy. BjCRFB1- or BjCRFB2-overexpressing cells were pretreated with RUX prior to GCRV infection. (M) Crystal-violet staining and absorbance quantification demonstrate that JAK inhibition completely abrogates BjCRFB-mediated cytoprotection and aggravates virus-triggered CPE. (N) qRT-PCR shows RUX markedly dampens BjCRFB-driven induction of downstream antiviral effector genes *viperin* and *Mx*. Mean ± SD; n = 3. Statistical significance: *\*P* < 0.05, *\*\*P* < 0.01, *\*\*\*P* < 0.001.

Considering the well-documented antiviral function of amphioxus Viperin^27^, we characterized the modulation of *viperin* transcription by BjSTATa and BjSTATb. At 4 h post viral infection, overexpression of either BjSTATa or BjSTATb significantly elevated *viperin* transcript abundance, with BjSTATb displaying markedly stronger transactivation capacity (Fig. 4H). We next generated a panel of truncated viperin promoter reporter constructs. Dual-luciferase reporter assays revealed that the P3 fragment (–1800 bp to +1) harbored the highest promoter activity, and this fragment was therefore chosen for further mechanistic validation. Subsequent co-transfection experiments demonstrated that both BjSTATa and BjSTATb robustly activated the transcriptional activity of the P3 promoter (Fig. 4I). We further predicted three potential STAT binding sites (BS1, BS2 and BS3) within the P3 region and generated site-specific mutant reporters. Mutations in BS2 or BS3 remarkably abolished BjSTATa/BjSTATb-mediated promoter activation (Fig. 4J). AlphaFold docking predicted three functional dimer configurations (BjSTATa–BjSTATa homodimer, BjSTATa–BjSTATb heterodimer, BjSTATb–BjSTATb homodimer), with BjSTATb homodimers exhibiting the highest promoter binding affinity (Supplementary Fig. S18). These findings demonstrate that BjSTATa and BjSTATb directly target the BS2 and BS3 cis-elements of the *viperin* promoter to initiate its transcription, uncovering the molecular mechanism underlying their antiviral function.

To genetically link BjCRFB receptor sensing to JAK–STAT transduction, we performed combinatorial pharmacological (ruxolitinib JAK inhibitor) and antibody blockade assays *in vivo* and *in vitro*. We first performed *in vivo* pharmacological blockade with the selective JAK inhibitor ruxolitinib (RUX). qRT-PCR on amphioxus gills showed that RUX pre-treatment drastically reduced antiviral *Mx* transcripts while elevating GCRV *VP5* viral loads (Fig. 4K), indicating intact JAK activity is required for tissue viral clearance. We next carried out combinatorial *in vivo* interference using BjCRFB1/2 neutralizing antibodies plus RUX co-administration. Dual suppression produced additive defects: *Mx* expression was further repressed, and *VP5* levels rose higher than single treatments alone (Fig. 4L). These data further corroborate that BjCRFB1 and BjCRFB2 function cooperatively with JAK kinases within antiviral signalling. Given potential compensatory immune circuits in whole animals, we validated this hierarchy via EPC cell gain-of-function assays. Crystal violet staining and quantitative absorbance measurements confirmed that JAK inhibition fully abolished the cytoprotective effects of BjCRFB1/2 overexpression and exacerbated virus-induced CPE (Fig. 4M). Consistently, qRT-PCR revealed RUX strongly attenuated BjCRFB1/2-driven induction of *Viperin* and *Mx* (Fig. 4N).

Overall, we demonstrate that amphioxus retains an undiversified, combinatorial ancestral JAK–STAT signalling module, establishing the evolutionarily conserved intracellular transduction logic shared by all chordate IFN cascades.

### BjMx acts as deeply conserved ancestral IFN-stimulated antiviral effector GTPase

Mx dynamin-like GTPases represent prototypical vertebrate ISGs with broad-spectrum antiviral activity^28–30^, but their ancestral form and function in basal chordates remained unvalidated. Phylogenetic analysis placed BjMx at the evolutionary boundary separating invertebrate and vertebrate Mx proteins (Supplementary Fig. S19). Despite low full-length primary sequence identity across taxa, BjMx harbours fully conserved GTP-binding domains and superimposable tertiary folds with human and zebrafish Mx (Fig. 5A, Supplementary Fig. S20, S21).

**Fig. 5:**
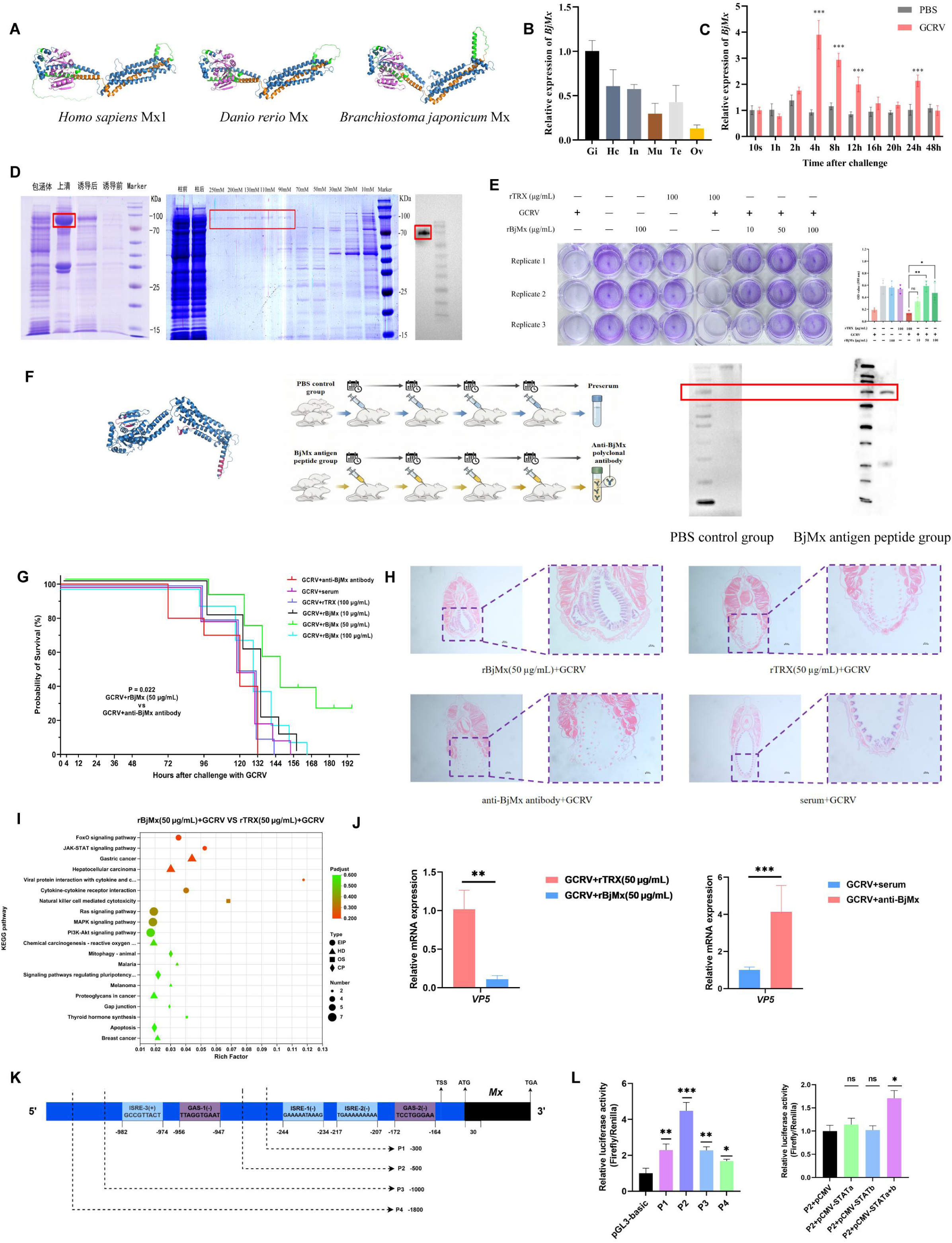
Amphioxus Mx represents an evolutionarily conserved IFN-stimulated antiviral effector. **(A)** Predicted three-dimensional structural models of Mx proteins from *Homo sapiens*, *Danio rerio* and *B. japonicum*. **(B)** qRT-PCR analysis showing tissue-specific expression profiles of *BjMx* in adult amphioxus. Gi, gill; Hc, hepatic caecum; In, hindgut; Mu, muscle; Te, testis; Ov, ovary. **(C)** Temporal expression patterns of *BjMx* in amphioxus following GCRV challenge, determined by qRT-PCR. **(D)** SDS–PAGE and western blot validation of purified recombinant His-tagged BjMx (rBjMx) produced in an *E. coli* expression system. **(E)** Crystal violet staining and quantitative cell viability assays showing dose-dependent protection conferred by rBjMx against GCRV infection in EPC cells. rTRX served as the negative control protein. **(F)** Schematic illustration of polyclonal antibody preparation against BjMx, and western blot validation confirming specific recognition of endogenous BjMx in amphioxus lysate by the custom anti-BjMx peptide antibody. **(G)** Kaplan–Meier survival curves of amphioxus subjected to GCRV challenge. Groups received supplementary rBjMx, rTRX control protein, or anti-BjMx neutralizing antibody to manipulate BjMx activity *in vivo*. **(H)** Representative H&E-stained histological sections of amphioxus hindgut from indicated treatment groups after GCRV infection, showing virus-triggered intestinal pathological lesions. **(I)** Bubble diagram summarizing significantly enriched KEGG pathways derived from comparative transcriptomic analysis of the rBjMx supplementation group versus the rTRX control group upon GCRV challenge. **(J)** qRT-PCR quantification of viral gene (*VP5*) transcript levels in amphioxus gills under gain-of-function (rBjMx treatment) and loss-of-function (anti-BjMx antibody neutralization) conditions after GCRV infection. **(K)** Schematic diagram of serially truncated BjMx promoter reporter constructs used for dual-luciferase assays. Key predicted cis-regulatory motifs are annotated along the promoter sequence. **(L)** Dual-luciferase reporter analysis defining the core active promoter fragment of BjMx (left), and luciferase assays evaluating the requirement of BjSTATa–BjSTATb heterodimerization for BjMx promoter activation (right). Data were from three independent experiments. All data represent means ± SD; statistical significance is indicated as \**P* < 0.05, \*\**p* < 0.01, \*\*\**p* < 0.001. ns, not significant.

Tissue expression analysis showed that *BjMx* is enriched in amphioxus immune tissues including gill, hepatic caecum and hindgut (Fig. 5B). GCRV infection significantly induced *BjMx* transcription from 4 h post-challenge (Fig. 5C), exhibiting typical ISG inducible expression patterns. We purified recombinant His-tagged rBjMx protein for functional assays (Fig. 5D). Exogenous rBjMx treatment conferred dose-dependent protection against GCRV in EPC cells (Fig. 5E). To further characterize the physiological role of endogenous BjMx *in vivo*, we generated a polyclonal antibody specific to BjMx, which efficiently recognizes native endogenous BjMx from *B. japonicum* (Fig. 5F). *In vivo* gain-of-function rBjMx supplementation and loss-of-function antibody neutralization assays showed that exogenous rBjMx increases amphioxus survival and attenuates virus-triggered intestinal damage. In contrast, antibody neutralization of endogenous BjMx accelerates host mortality and exacerbates pathological lesions (Fig. 5G, H).

Comparative transcriptomic analysis of rBjMx gain-of-function and antibody-mediated loss-of-function models revealed BjMx-dependent immune regulatory networks. rBjMx supplementation during GCRV infection activated core innate immune pathways, including JAK–STAT antiviral signalling, FoxO-mediated oxidative stress homeostasis and cytokine receptor interactions (Fig. 5I), while effectively restricting viral replication(Fig. 5J). In contrast, BjMx depletion triggered inflammatory lipid metabolism dysregulation and compensatory broad-spectrum antiviral responses (Supplementary Fig. S22), accompanied by significantly elevated viral loads (Fig. 5J). These data demonstrate that BjMx remodels host antiviral transcriptional profiles and maintains immune homeostasis during viral infection.

To further probe the transcriptional regulatory mechanism governing Mx expression in amphioxus, we conducted serial truncation and dual-luciferase reporter assays to map the core active region of the BjMx promoter. Promoter truncation and dual-luciferase assays identified core cis-regulatory elements within the -500 to -300 bp region of the *BjMx* promoter (Fig. 5K). Distinct from *Viperin* regulation, *BjMx* promoter activation strictly requires BjSTATa–BjSTATb heterodimerization (Fig. 5L), revealing early differentiated effector regulatory circuits within the primordial IFN network.

## Discussion

### The complete IFN antiviral signalling network evolved in basal cephalochordates prior to vertebrate radiation

Decades of comparative immunological research have outlined stark evolutionary divides in antiviral innate immunity across metazoan lineages. Protostome groups including molluscs and arthropods only carry partial components of the JAK–STAT axis, lacking canonical IFN ligands and dedicated IFN receptors entirely. Their TBK-related kinases lack any capacity for immune signal transduction and instead function in developmental regulation; coupled with the absence of IRF transcription factors that abrogates their ability to launch IFN-dependent transcriptional programmes, these invertebrates are incapable of mounting authentic IFN-mediated antiviral defence^1,7,31^. Cross-kingdom comparative analyses of antiviral defence have recently unified prokaryotic anti-phage systems and eukaryotic innate immunity into a single evolutionary framework, revealing that core nucleic acid sensing, signal amplification and effector modules trace their origin to ancient bacterial immune machineries, yet protostome lineages failed to retain the full set of building blocks required for IFN cascades spanning viral detection to effector execution^32^. By contrast, jawed vertebrates such as teleosts and mammals possess elaborate multi-layered IFN cascades featuring diversified IFN subtypes, specialized receptor assemblies and a broad panel of interferon-stimulated antiviral effectors that govern tissue-specific immune homeostasis against diverse pathogens^3,7^. Comprehensive reviews centred on interferon physiology and pathology have systematically summarized the layered, tissue-differentiated functions of type I, II and III IFNs across vertebrate species, highlighting how subtype specialization balances robust viral clearance and controlled inflammatory responses to avoid host tissue damage^33^. This long-noted phylogenetic separation has sustained a prevailing evolutionary model holding that fully functional IFN signalling originated exclusively within vertebrate lineages(Fig.6).

**Fig. 6.**
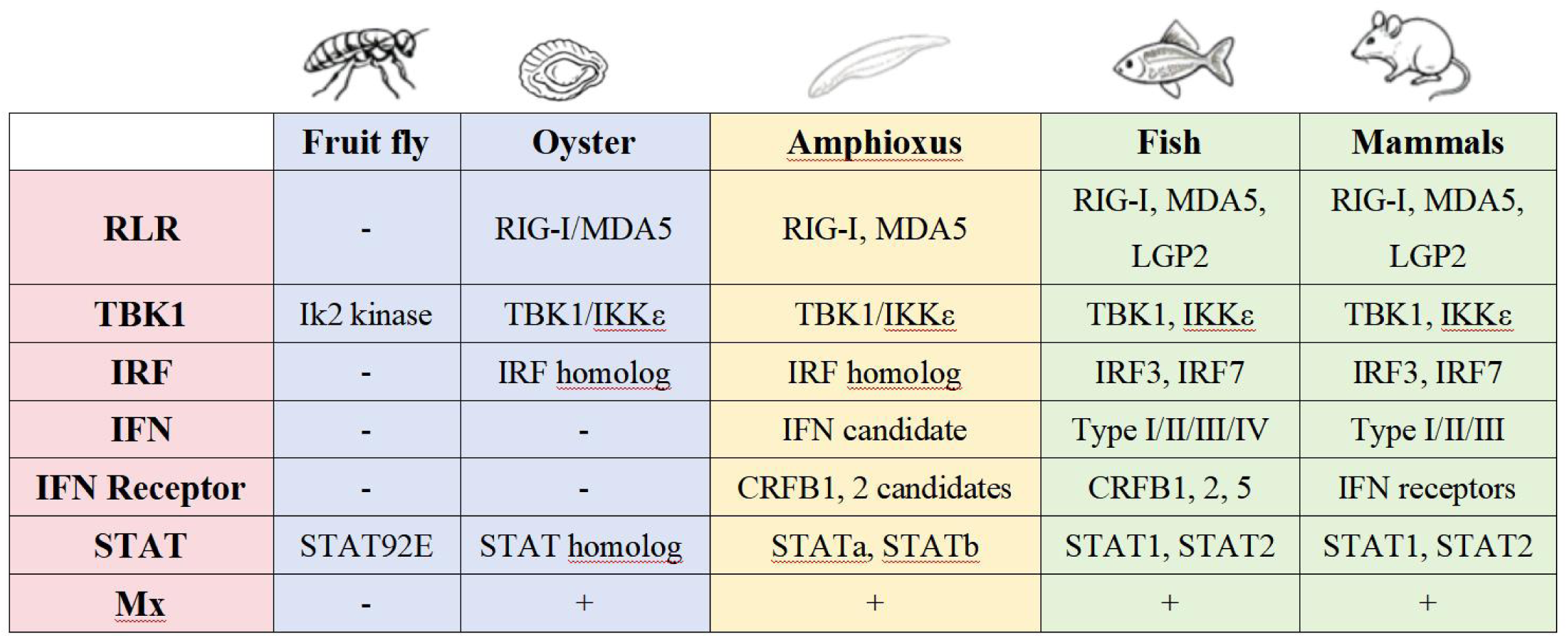
Evolutionary conservation of core antiviral signaling components across metazoans. Comparison of key IFN-pathway molecules among fruit fly, oyster, amphioxus, fish and mammals. Symbols “−” and “+” indicate gene absence and presence, respectively. Homologs represent sequence-identified family members, and “candidate” denotes amphioxus primordial IFN-related molecules characterised in the present work. This cross-species comparison reveals that amphioxus already harbours the core genetic toolkit for an ancestral IFN-like antiviral network prior to the emergence of vertebrate canonical IFN systems.

Our integrated structural, phylogenetic and *in vitro* plus *in vivo* functional characterization of the complete IFN cascade in *B. japonicum* revises this paradigm entirely. The full hierarchical signalling circuit we reconstitute in amphioxus encompasses every core module present in vertebrate IFN pathways, ranging from TBK1/IKKε-dependent IFN transcription induction and cell-surface sensing by primitive BjCRFB receptors to canonical intracellular JAK–STAT transduction and terminal restriction mediated by conserved Mx and Viperin effector proteins (Fig.7). As the sister clade to all vertebrates, cephalochordate amphioxus occupies a pivotal transitional evolutionary position bridging simple invertebrate innate defence and sophisticated vertebrate IFN immunity, providing definitive functional proof that IFN-dependent antiviral signalling constitutes an ancestral chordate trait rather than a vertebrate evolutionary novelty.

**Fig. 7:**
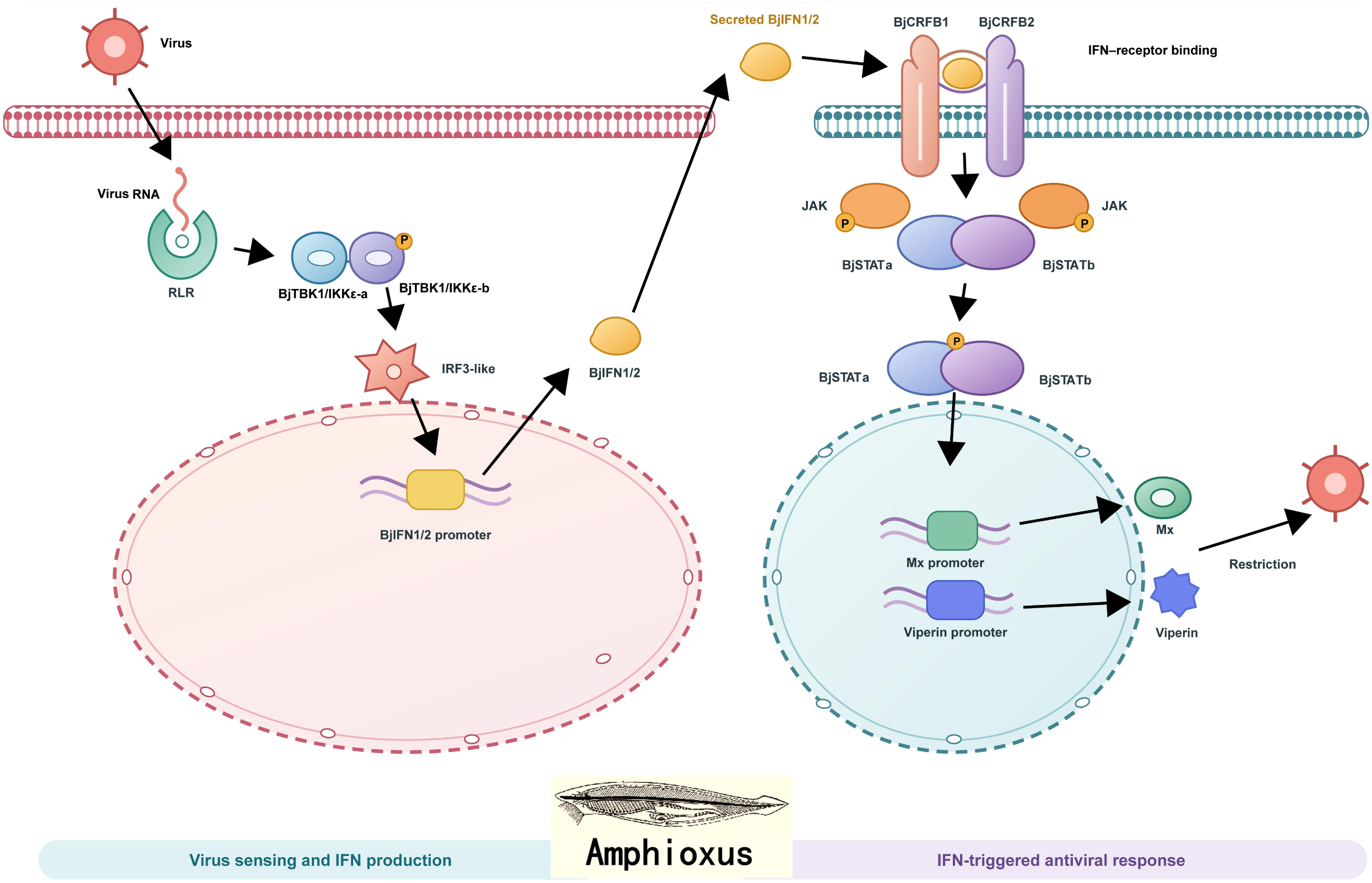
Working model for the primordial IFN antiviral signalling network in amphioxus. A schematic representation of the ancestral IFN-dependent antiviral pathway uncovered in *B. japonicum*. Left panel: viral nucleic acids are detected via RLR sensing, leading to BjTBK1/IKKε-mediated activation of IRF3-like factors and subsequent transcription of BjIFN1/2. Secreted BjIFN1/2 binds the BjCRFB1-BjCRFB2 receptor complex (right panel), initiating JAK-STAT signalling. Phosphorylated BjSTATa/BjSTATb dimers enter the nucleus to induce antiviral effector genes such as *Mx* and *Viperin*, which mediate viral restriction. This model depicts a pre-vertebrate prototype of the canonical vertebrate IFN antiviral system.

### Ancestral molecular signatures of amphioxus IFN cascade trace stepwise evolutionary refinement of chordate antiviral immunity

Multidimensional functional profiling of the amphioxus IFN pathway reveals a suite of plesiomorphic molecular features that collectively trace the incremental optimization of chordate antiviral defence. Primordial IFN biological activity is maintained primarily through conserved compact α-helical tertiary folds rather than rigid linear amino acid sequences. Persistent pathogen-driven selective pressure across vertebrate radiation has driven extensive divergence in IFN primary sequences, while the helical core architecture indispensable for ligand function has remained under strict structural constraint. This stable conserved folding framework laid the molecular foundation for successive rounds of gene duplication and paralog specialization in vertebrates, giving rise to tissue-specific IFN subtypes tailored to distinct infectious challenges, a layered functional differentiation thoroughly documented in vertebrate interferon immunology^33^. Another defining ancestral characteristic lies in the weak, transient binding kinetics observed between amphioxus IFN ligands and their cognate BjCRFB receptors. All AlphaFold3 structural docking predictions for BjIFN–BjCRFB complexes yield low interface confidence metrics, reflecting loose, evolutionarily primitive interaction geometry that stands apart from the high-affinity, stable ternary receptor assemblies found in mammalian and teleost IFN systems. The enhancement of ligand–receptor binding affinity represents a vertebrate-specific adaptive advance that elevates the sensitivity of pathogen surveillance within complex microenvironments rich in diverse microbes and viruses. Amphioxus also retains flexible, partially overlapping functional profiles for duplicated TBK and STAT signalling mediators, consistent with universal defensive strategies shared across bacteria and eukaryotes, where undiversified multi-functional signalling complexes represent the ancestral immune state before lineage-specific subfunctionalization^32^. Vertebrate evolution later partitioned discrete signalling tasks among duplicated kinase and STAT paralogs via gene duplication and subfunctionalization, enabling precise, stimulus-tailored transcriptional outputs upon viral exposure. Taken together, these ancestral molecular traits support a unified evolutionary trajectory for chordate IFN immunity: basal cephalochordates assembled the full core architectural framework required for canonical IFN signalling, and vertebrate lineages progressively refined this primitive immune toolkit through accumulated sequence variation, strengthened ligand–receptor interactions and the compartmentalization of duplicated signalling mediators.

Phylogenetic and tertiary structural analyses delineate the evolutionary origin of amphioxus primitive IFN receptors BjCRFB1 and BjCRFB2. Phylogenetic trees built from extracellular domain sequences place both proteins within the class II cytokine receptor superfamily, yet neither establishes direct one-to-one orthology to any vertebrate IFN receptor subtype. Structural superimposition against human and teleost receptors reveals differential fold similarity: BjCRFB2 shares higher TM-score homology with human IFNGR2 and zebrafish type I IFN co-receptor CRFB5, whereas BjCRFB1 exhibits moderate structural resemblance to human IL10RA. Conserved FN3 domain alignments uncover no subtype-specific signature motifs to categorize either BjCRFB paralog into vertebrate type I, II or III IFN receptor groups. Integrating phylogenetic and structural evidence, BjCRFB1 and BjCRFB2 constitute undifferentiated ancestral receptor prototypes arising before vertebrate IFN receptor subfunctionalization. While sequence and fold data cannot confirm strict orthology, their virus-triggered expression kinetics echo vertebrate IFNLR receptors specialized for mucosal antiviral defence^33^. BjCRFB2 mediates rapid acute immune signalling, and BjCRFB1 sustains prolonged antiviral responses—this temporal dichotomy mirrors the long-term epithelial surveillance governed by IFN-λ receptors, suggesting a tentative functional connection to ancestral type III IFN pathways.

Combined structural modelling, docking predictions and tissue expression profiles support tentative evolutionary inference linking BjIFN1/2 to vertebrate IFN subtypes. Amphioxus IFNs share negligible primary sequence identity with vertebrate counterparts, driven by persistent pathogen-mediated diversifying selection conserved across all cellular immune lineages. Despite poor linear homology, chordate IFNs are unified by a compact α-helical scaffold critical for ligand function. BjIFN1 and BjIFN2 lack the extended C-terminal helical segments of vertebrate type I IFNs^34^. Their compact helical core hints at potential structural resemblance to mucosa-specialized vertebrate IFN-λ^33,34^, which remains an unvalidated hypothesis. This proposed type III-like ancestral identity of amphioxus IFNs illuminates the evolutionary origins of chordate mucosal antiviral immunity. Transcriptomic and in situ hybridization data show BjIFNs and their BjCRFB receptors concentrate in gill and hindgut epithelia, the primary pathogen-exposed barriers of filter-feeding amphioxus. As summarized in vertebrate interferon reviews^33^, type III IFNs act as localized mucosal sentinels to restrict viral spread without systemic inflammation, limiting organism-wide immunotoxicity—this tissue-specific immune strategy first arose in basal chordates via the primitive BjIFN–BjCRFB signalling axis. Weak ligand-receptor affinity acts as an adaptive brake to avoid aberrant systemic activation under constant low aquatic pathogen exposure. Vertebrates later refined this ancestral mucosal IFN framework via serial gene duplication, enhanced binding avidity and the evolution of systemic type I IFN cascades, establishing a dual mucosal-systemic antiviral network derived from cephalochordate primordial immunity.

### Deep evolutionary conservation and incipient subfunctionalization across all IFN cascade modules

Each component of the amphioxus IFN pathway retains deeply conserved functional properties across chordate lineages while displaying early-stage subfunctionalization that laid the groundwork for the diversified vertebrate IFN toolkit. The upstream TBK1/IKKε–IRF regulatory axis stands as the most evolutionarily invariant segment of the entire signalling cascade. Conserved TBK-IRF nucleic acid sensing modules are widely distributed across prokaryotic anti-phage defence and eukaryotic innate immune systems as core ancient immune machinery^40^. Amphioxus BjTBK1/IKKε-a and BjTBK1/IKKε-b are undifferentiated ancestral precursors predating vertebrate TBK1/IKKε subfamily divergence; their cytoplasmic localization, heterodimerization capacity and phosphorylation-dependent activation switches are fully conserved with mammalian and teleost homologs. Amphioxus IRF3-like transcription factors retain the core function of driving IFN ligand transcription, confirming that the TBK– IRF-dependent IFN production circuit originated in basal chordates and remained highly conserved throughout chordate evolution. The BjCRFB receptor module balances structural conservation with early functional divergence. Shared FN3 domain architecture and virus-inducible antiviral activity confirm that vertebrate IFN receptors evolved from ancestral CRFB-like precursors in cephalochordates, while divergent temporal response dynamics and differential structural affinity toward distinct vertebrate receptor clades mark the initial stage of immune sensor specialization, laying the evolutionary groundwork for the broad spectrum of IFN receptor subtypes seen in modern vertebrates. The JAK–STAT transduction cascade and terminal Mx effector GTPase display near-perfect functional conservation across all chordate lineages. The STAT dimer-dependent transcriptional activation mechanism and strict JAK kinase dependence are identical between amphioxus and vertebrates; BjMx retains conserved tissue enrichment, virus-inducible expression and broad-spectrum GTPase-mediated antiviral activity characteristic of vertebrate ISG Mx proteins. Notably, differential STAT dimer requirements for Viperin versus Mx promoter activation reveal early effector regulatory divergence within the primordial IFN network, a trait further amplified and refined during vertebrate IFN system diversification. Collectively, patterns of deep conservation paired with incipient paralog differentiation across all signalling tiers confirm that the full set of core IFN signalling modules was already assembled in the most basal extant chordates, and vertebrate IFN cascades represent diversified, optimized derivatives inherited from this cephalochordate ancestral toolkit rather than independent evolutionary innovations.

### Broader evolutionary implications for comparative innate immunology

Our findings rewrite the established evolutionary timeline of chordate IFN immunity. Prior comparative genomic and functional studies assumed complete, functional IFN signalling arose within vertebrate lineages. In contrast, our multi-layered structural and *in vivo* functional evidence definitively places the origin of the full IFN antiviral network at the cephalochordate grade, demonstrating IFN-mediated innate immunity as an ancestral chordate synapomorphy. Primitive molecular hallmarks observed in amphioxus—low primary sequence but high structural conservation of IFN ligands, weak ligand-receptor binding interactions, and flexible undifferentiated STAT signalling modules—represent an adaptive immune mode suited to the low pathogen complexity of early chordate aquatic habitats. As vertebrates diversified and pathogen pressure intensified across terrestrial and aquatic niches, cumulative sequence mutation, optimized molecular binding affinity and paralog functional compartmentalization progressively refined IFN signalling into the efficient, stimulus-specific antiviral cascades observed in modern vertebrates. Cross-kingdom comparative frameworks laid out in recent work unifying human and bacterial antiviral immunity provide a global evolutionary context for our observations^40^: core sensing, amplification and effector machineries originated from bacterial anti-phage defence systems, and chordates uniquely assembled these ancient building blocks into a complete IFN-specific antiviral cascade absent in all protostome invertebrate taxa, while vertebrate-specific subtype expansion and tissue specialization further fine-tuned IFN function for complex multicellular immune homeostasis^41^.

### Study limitations and future perspectives

While this work systematically reconstructs and validates the complete primordial IFN signalling cascade in *B. japonicum*, several critical mechanistic and evolutionary questions remain unaddressed and require further investigation. First, the upstream adaptor proteins bridging cytoplasmic RLR/TLR pattern recognition receptors and the BjTBK1/IKKε heterocomplex have not yet been identified, leaving an uncharacterized gap at the uppermost tier of viral signal transduction; universal cross-kingdom immune logic indicates such bridging adaptors are a core component of ancient nucleic acid sensing circuits^32^, and targeted interactome screening will be necessary to fill this missing signalling link. Second, our interpretations of BjSTATa and BjSTATb functional divergence are constrained by single timepoint sampling; time-resolved viral stimulation gradients are required to disentangle time-layered transcriptional responses from intrinsic paralog specialisation. Third, high-resolution cryo-electron microscopy combined with surface plasmon resonance biochemical assays will be essential to resolve atomic-level differences in ligand-receptor recognition and STAT dimer DNA binding affinity. Fourth, the full pathogen response spectrum of the amphioxus primordial IFN network remains poorly defined, and its molecular crosstalk with complement, inflammatory signalling and autophagy regulatory circuits requires systematic multi-omic dissection.

Moving forward, integrated multi-omics profiling, high-resolution structural immunology and comparative functional assays across diverse basal chordate species will help fully resolve the stepwise structural and functional fine-tuning of IFN immunity accumulated over hundreds of millions of years of chordate diversification, and further bridge evolutionary links between prokaryotic anti-phage defence and vertebrate interferon networks^32^.

## Conclusion

Here, we report the full functional reconstitution of an intact primordial IFN antiviral signalling network in cephalochordate amphioxus, the most basal extant chordate lineage. This ancestral signalling cascade covers TBK1/IKKε–IRF-mediated virus-dependent IFN transcription, primitive BjCRFB cell-surface IFN receptors, canonical JAK–STAT intracellular signal transduction, and conserved Mx/Viperin antiviral effector proteins. The amphioxus IFN system retains hallmark ancestral plesiomorphic traits. These features include structural conservation of IFN ligands, primitive weak binding affinity between ligands and receptors, and undiversified flexible signalling transducers. Together these molecular signatures outline a gradual trajectory of structural and functional refinement for vertebrate innate IFN immunity. Our findings definitively trace the evolutionary origin of vertebrate interferon-mediated antiviral defence to basal cephalochordates, resolve a decades-long major gap in evolutionary immunology, and establish a comprehensive conceptual framework to understand the structural diversification and functional optimization of innate antiviral signalling across all chordate lineages.

## Methods

### Animals and virus strains

Laboratory-bred *Branchiostoma japonicum* adults were maintained in filtered natural seawater at 22 ± 1 °C under standard culture conditions. Genotype I grass carp reovirus (GCRV) was provided by Prof. Yibing Zhang (Institute of Hydrobiology, CAS). Viral titres were quantified via TCID_50_ assay using the Reed–Muench method^35^. All amphioxus experiments followed ethical guidelines approved by the Animal Protection and Use Committee of Ocean University of China (Permit SD2007695).

### Cell lines and cell culture

EPC (Epithelioma Papulosum Cyprini) cells were acquired from the China Zebrafish Resource Center (CZRC). Human HEK293T cells were obtained from ATCC. EPC cells were cultured in MEM medium supplemented with 10% FBS, 100 U/mL penicillin and 100 μg/mL streptomycin at 28 °C, 5% CO₂. HEK293T cells were cultured in DMEM with 10% FBS at 37 °C, 5% CO₂.

#### Gene cloning and plasmid construction

Total RNA was extracted from amphioxus tissues using TRIzol reagent, reverse-transcribed into first-strand cDNA. Target ORFs were amplified via gene-specific primers. Purified PCR fragments were subcloned into pCMV-C-Flag, pcDNA3.1-EGFP, pGL3-basic and pET-28a vectors for overexpression, subcellular localization, dual-luciferase reporter and prokaryotic recombinant protein expression, respectively. Site-directed mutagenesis was performed using commercial kits, and all plasmids were verified by Sanger sequencing.

### Bioinformatic and structural analyses

Signal peptide prediction (SignalP 6.0)^36^, transmembrane domain prediction (TMHMM v2.0)^37^, domain annotation (SMART, Pfam)^38,39^, multiple sequence alignment (MAFFT)^40^ and maximum-likelihood phylogenetic trees (MEGA11, 1000 bootstrap replicates)^41^ followed standard protocols. Protein tertiary structures and ligand–receptor docking complexes were predicted with AlphaFold3^42^; global structural comparison was performed using TM-align^43^, for which ipTM scores and inter-chain predicted aligned error (PAE) were used to assess confidence in the predicted inter-chain geometry.

### *In vitro* viral infection

EPC cells were seeded 24 h pre-transfection, challenged with GCRV (10⁵ TCID50/mL) in serum-free medium for 1 h, washed and replenished with complete medium. Cell samples were harvested at designated timepoints for qPCR, CCK-8 viability, crystal violet staining and western blot assays.

### *In vivo* GCRV challenge

Adult amphioxus were fasted for 24 h before viral challenge. For each biological replicate, 10 amphioxus were maintained in 200 mL of sterile seawater, after which 40 mL of GCRV-Amphi2025 suspension at a titre of 10⁵ TCID₅₀/mL was added to the infection group. Control groups received equivalent cell culture medium. Survival curves (Kaplan– Meier) and tissue RNA collection were performed at defined timepoints; each biological group contained 10 individuals with three independent replicates.

### Tissue histology

Dissected gill and hindgut tissues were fixed in 4% PFA for 24 h, paraffin-embedded, sectioned to 5 μm thickness and stained with haematoxylin and eosin (H&E). Tissue lesion area was quantified via ImageJ (v1.54q) morphometric analysis.

### Transfection and cellular functional assays

HEK293T and EPC cells used in the paper were transfected with Lipofectamine 3000 (Invitrogen) according to the manufacturer’s instructions. Subcellular localization was visualized via laser scanning confocal microscopy. CCK-8 and crystal violet staining quantified cell viability and cytopathic damage. Dual-luciferase reporter activity was detected using Luc-Pair Duo-Luciferase Kit 2.0, with firefly luciferase normalized to Renilla internal control.

### Co-immunoprecipitation and western blot

Cell lysates were incubated with anti-Flag/anti-HA conjugated beads overnight at 4 °C. Eluted protein complexes were separated by 12.5% SDS–PAGE, transferred to PVDF membranes and blocked with 5% non-fat milk. Primary antibodies (anti-Actin, anti-HA, anti-Flag, anti-His) and HRP-conjugated secondary antibodies were used for ECL chemiluminescence detection.

### Polyclonal antibody production

Peptides corresponding to extracellular domains of BjCRFB1, BjCRFB2 and full-length BjMx were selected for antigen synthesis, conjugated to KLH carrier protein and injected into rats for immunization. Crude antiserum was purified via peptide affinity chromatography; antibody specificity was validated by western blot prior to in vivo neutralization experiments.

### *In vivo* antibody and pharmacological blockade

Purified anti-BjCRFB1, anti-BjCRFB2 and anti-BjMx polyclonal antibodies were microinjected into amphioxus for loss-of-function assays. For JAK inhibition, ruxolitinib was pre-administered prior to GCRV challenge. Kaplan–Meier survival analysis, gill tissue viral load and ISG qPCR quantification were performed for all treatment groups.

### Transcriptome sequencing and enrichment analysis

Total RNA from rBjMx supplementation, anti-BjMx and antiCRFB1/2 neutralization groups was extracted for Illumina HiSeq 4000 sequencing by Annoroad Technology (Beijing, China) (three biological replicates per group). Differentially expressed genes (fold change ≥2, FDR < 0.05) were identified using edgeR; KEGG pathway enrichment analysis was performed to characterize perturbed immune signalling networks.

### Quantitative real-time PCR (qRT-PCR)

Total RNA was DNase-treated and reverse-transcribed; Gene expression was determined by amplifying the cDNA with ChamQ SYBR Color qPCR Master Mix(Vazyme, #Q431-02) by an ABI 7500 Fast Real-Time PCR System (Applied Biosystems). The 2^−ΔΔCt^ method was used to calculate relative expression changes, with amphioxus *ef-1ɑ* or EPC *actin* as internal reference. All reactions were run in technical triplicates across three biological replicates.

### Statistical analysis

All experiments were repeated at least three times independently. All statistical analysis was performed using Graphpad Prism 8.0.2. Data are presented as mean ± SD. Statistical significance between two groups was determined by two-tailed student’s *t*-test. For comparison of multiple groups, the statistical analysis was performed using a one-way ANOVA followed by Games-Howell post hoc tests. Survival analyses were performed using the Kaplan-Meier method and assessed using the log-rank (Mantel-Cox) test. \**p* < 0.05, \*\**p* < 0.01, *** *p*< 0.001; ns, not significant, *p* > 0.05.

## Data availability

All raw sequencing datasets, structural prediction files, source imaging data and primer sequences are available from the corresponding author upon reasonable written request. GenBank accessions for amphioxus IFN paralogs: PZ746091 (BjIFN1), PZ746092 (BjIFN2). All supplementary tables and figure source data are provided with this manuscript submission.

## Author contributions

Z.-H. L., G.-D. J. and C. S. conceived and designed the research project and coordinated the experimental implementation. J.-Y. L. performed the experiments and analyzed the data for the amphioxus IFN and CRFB. Y.-L. H. performed the experiments and analyzed the data for the amphioxus TBK1/IKK. H.-J. Y. performed the experiments and analyzed the data for the amphioxus STAT. M. Y. performed the experiments and analyzed the data for the amphioxus Mx. Z.-H. L. and J.-Y. L. drafted and revised the manuscript. All authors reviewed and approved the final version of the manuscript.

## Acknowledgements

This work was funded by the grants of National Natural Science Foundation of China (grant number: 32570617) and Science & Technology Innovation Project of Laoshan Laboratory (grant number: LSKJ202203204). We sincerely thank Shenjie Zhong for his contribution to the identification of amphioxus CRFB receptors. We also thank Jiayi Shi and Rongxiao Zhang for analysis of amphioxus IRF.

## Conflict of Interest

The authors declare that they have no conflict of interest.

## Notes

### Competing Interest Statement

The authors have declared no competing interest.

