## Supplementary Tables and Figures for "An intact primordial interferon antiviral signalling network in amphioxus illuminates the basal chordate origin of vertebrate IFN immunity"

**Table S1**: Comparison of basic information and key primary-structural parameters of six amphioxus IFN-like proteins

| **Protein** | **Species** | **ID** | **Length (aa)** | **MW (kDa)** | **Theoretical pI** | **Predicted N-glyco motifs** |
|---|---|---|---|---|---|---|
| BfIFN1 | *B. floridae* | PZ236457 | 207 | 22.79 | 5.43 | 0 |
| BfIFN2 | *B. floridae* | PZ220848 | 215 | 23.69 | 8.25 | 0 |
| BbeIFN1 | *B. belcheri* | XM_019780013 | 207 | 23.08 | 8.60 | 0 |
| BbeIFN2 | B. belcheri | PZ220849 | 207 | 23.08 | 8.85 | 0 |
| BjIFN-like-A | *B. japonicum* | PZ746091 | 208 | 22.99 | 8.25 | 0 |
| BjIFN-like-B | *B. japonicum* | PZ746092 | 207 | 22.87 | 7.60 | 1 |

**Table S2**: TM-score Statistics for Three-dimensional Structural Similarity between BjCRFB1 and Human Class II Cytokine Receptors

| Target | TM-score |
|---|---|
| *Homo sapiens* IFNAR1 | 0.37771 |
| *Homo sapiens* IFNAR2 | 0.34430 |
| *Homo sapiens* IFNGR1 | 0.34632 |
| *Homo sapiens* IFNGR2 | 0.35070 |
| *Homo sapiens* IFNLR1 | 0.37577 |
| *Homo sapiens* IL10RA | 0.40321 |
| *Homo sapiens* IL10RB | 0.38369 |

**Table S3**: TM-score Statistics for Three-dimensional Structural Similarity between BjCRFB2 and Human Class II Cytokine Receptors

| Target | TM-score |
|---|---|
| *Homo sapiens* IFNAR1 | 0.42869 |
| *Homo sapiens* IFNAR2 | 0.45509 |
| *Homo sapiens* IFNGR1 | 0.51968 |
| *Homo sapiens* IFNGR2 | 0.53364 |
| *Homo sapiens* IFNLR1 | 0.48677 |
| *Homo sapiens* IL10RA | 0.47151 |
| *Homo sapiens* IL10RB | 0.49677 |

### Supplementary Figures

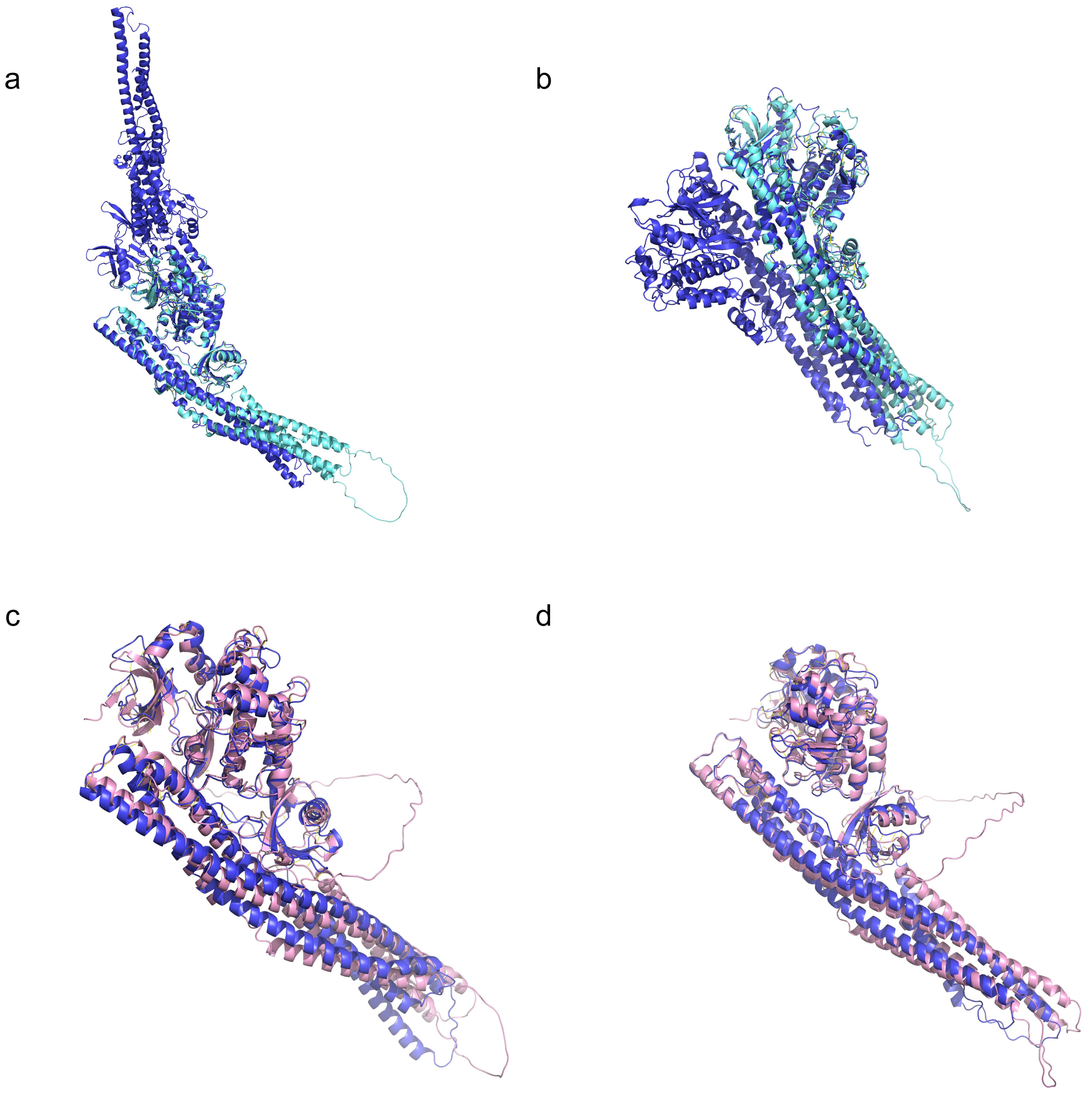


**Fig.S01**: **Cross-species structural comparison reveals high evolutionary conservation of BjTBK1/IKKε-a and BjTBK1/IKKε-b**. Structural superimposition of amphioxus BjTBK1/IKKε paralogs against vertebrate TBK1 and IKKε orthologs. **a:** superimposition of zebrafish (*Danio rerio*) DrTBK1 with BjTBK1/IKKε-a (blue). **b:** superimposition of human (*Homo sapiens*) HsTBK1 with BjTBK1/IKKε-a (blue). **c:** superimposition of zebrafish DrIKKε with BjTBK1/IKKε-b (pink). **d:** superimposition of human HsIKKε with BjTBK1/IKKε-b (pink). Amphioxus BjTBK1/IKKε-a and BjTBK1/IKKε-b display comparable tertiary-structure homology to both vertebrate TBK1 and IKKε, without preferential similarity toward either vertebrate subfamily.

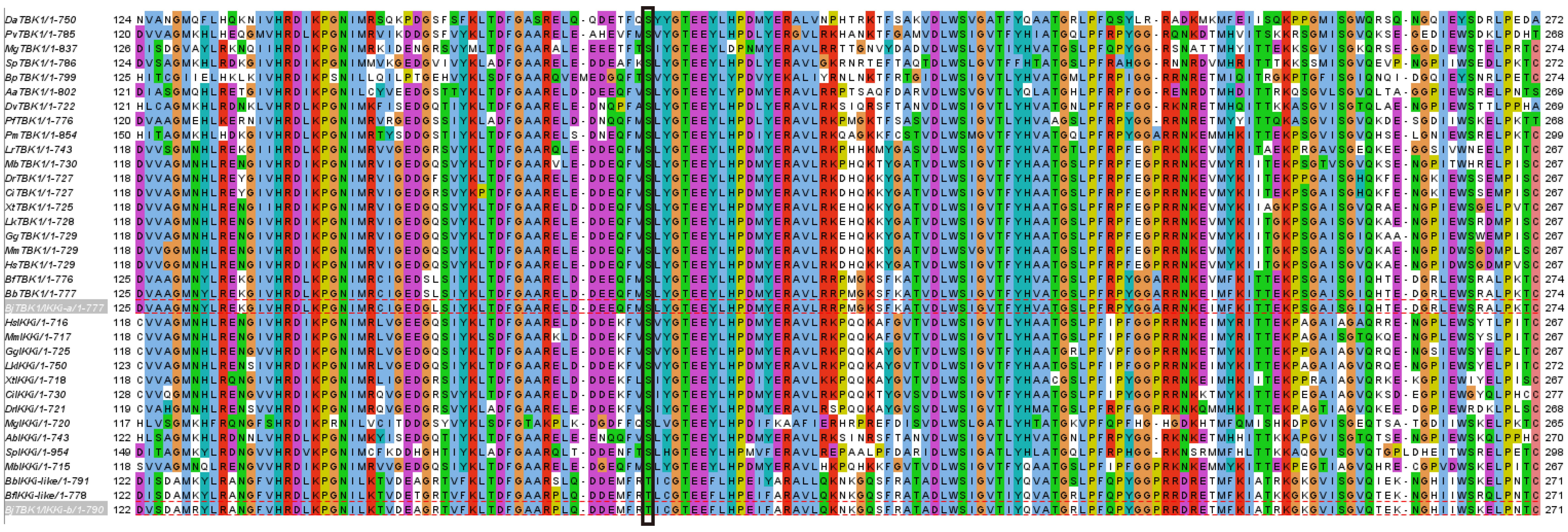

**Fig.S02: Multiple sequence alignment of the kinase domain (KD) from TBK1/IKKε family proteins across metazoan species.** Multiple sequence alignment of the kinase domain region of TBK1/IKKε homologs from diverse animal lineages. Tthe serine/threonine phosphorylation residues: Ser179 in BjTBK1/IKKε-a and Thr176 in BjTBK1/IKKε-b are broadly conserved among TBK1/IKKε orthologs across vertebrate and invertebrate species.

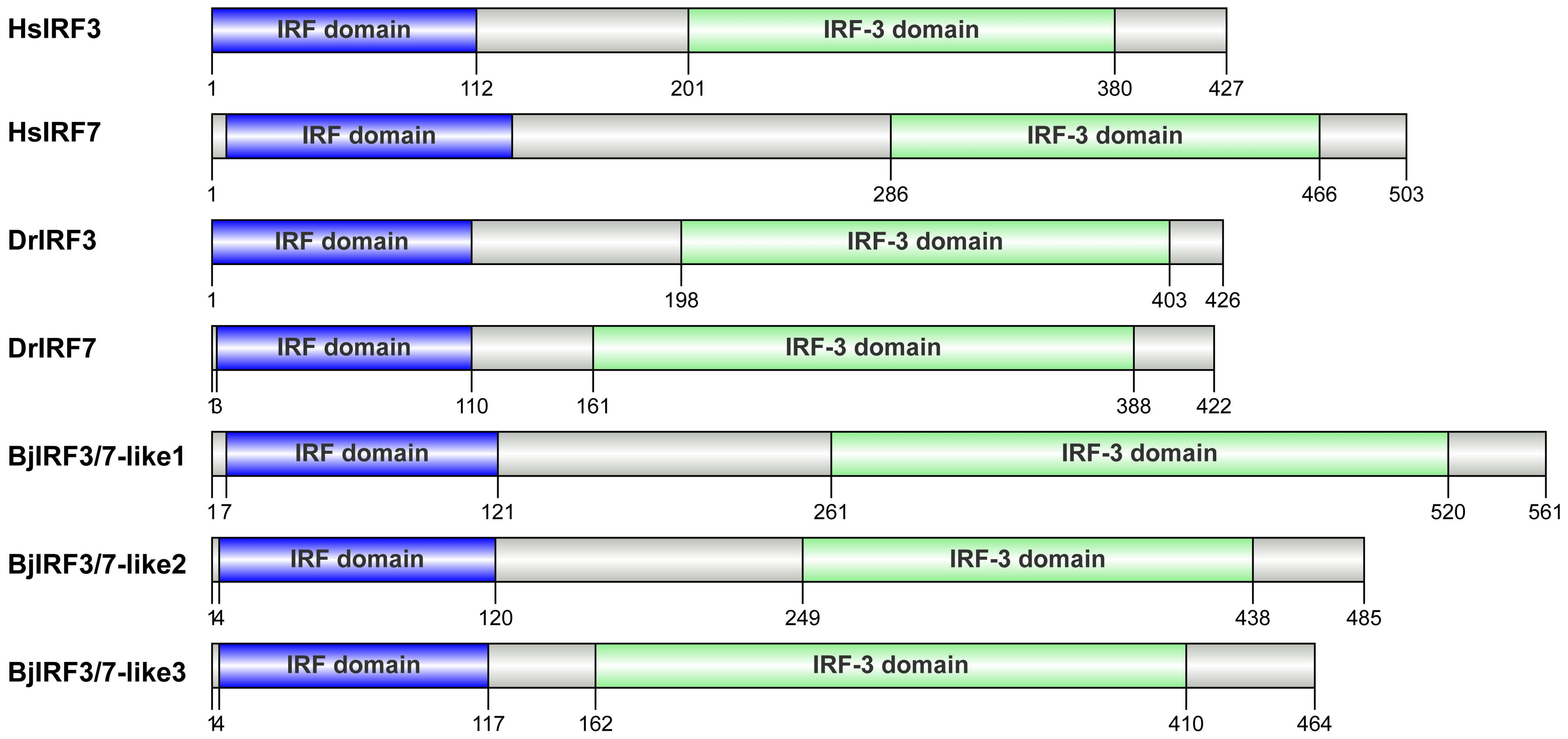


**Fig.S03: Domain architecture of amphioxus IRF family proteins.** Schematic domain organization of multiple IRF family members identified from *Branchiostoma japonicum* genome. Purple boxes denote the interferon regulatory factor transcription factor domain; green boxes represent the conserved IRF3-specific functional domain. Several amphioxus IRF proteins possess intact IRF3-related functional domains, supporting their potential roles as ancient IRF3-like transcription factors. Related phylogenetic analysis is shown in Figure S04.

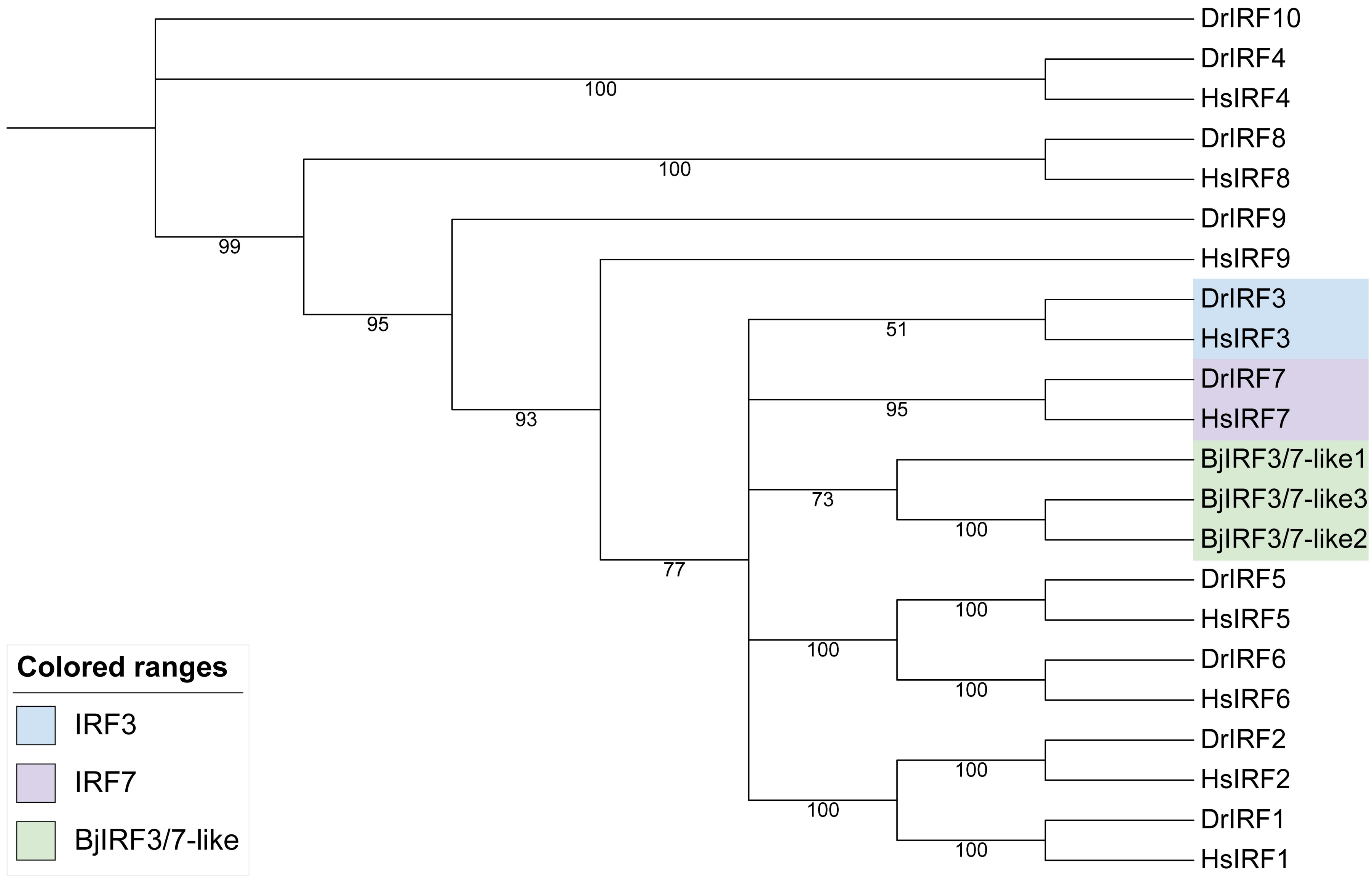


**Fig.S04: Phylogenetic tree of IRF family proteins from vertebrates and amphioxus.** Maximum-likelihood phylogenetic reconstruction of IRF family members from human (*Homo sapiens*, Hs), zebrafish (*Danio rerio*, Dr), and *Branchiostoma japonicum* (Bj). Numbers at internal nodes denote bootstrap support values. Colour shading: light-blue, vertebrate IRF3 clade; light-purple, vertebrate IRF7 clade; light-green, amphioxus BjIRF-like candidates carrying IRF3-type domains identified in Fig. S03. Amphioxus IRF candidates group within the same large clade containing vertebrate IRF3/IRF7, indicating close evolutionary relationship to vertebrate IRF3 homologs.

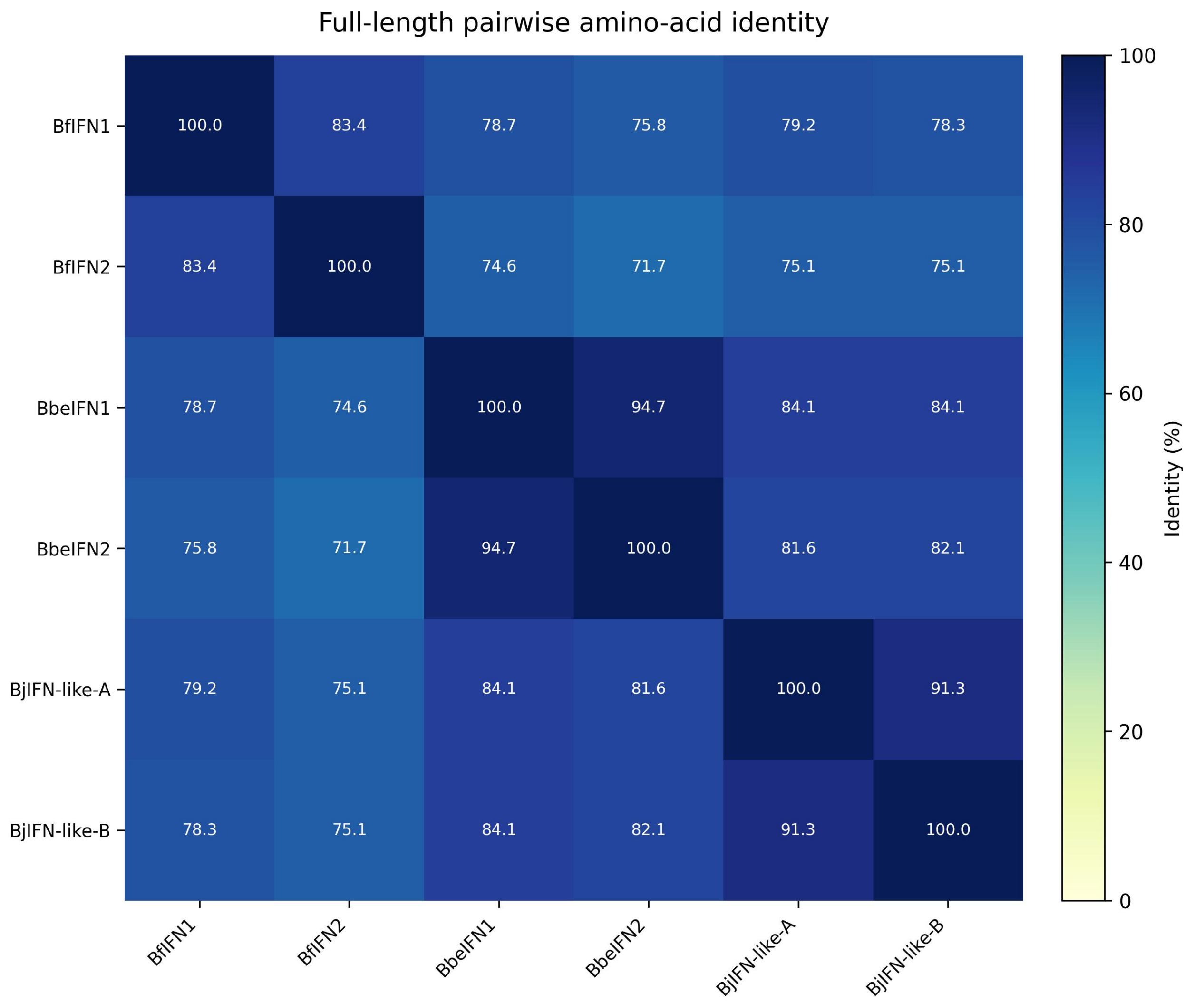


**Fig.S05**: **Full-length pairwise amino-acid identity among six amphioxus IFN-like proteins.** Heatmap displaying pairwise amino acid sequence identity (%) across full-length sequences of six amphioxus IFN-like proteins, including *Branchiostoma floridae* BfIFN1, BfIFN2, *Branchiostoma belcheri* BbeIFN1, BbeIFN2 and *Branchiostoma japonicum* BjIFN-like-A (BjIFN1), BjIFN-like-B (BjIFN2). The colour gradient bar represents sequence identity from 0% (light yellow) to 100% (dark navy-blue). Percent identity values are shown within each cell.

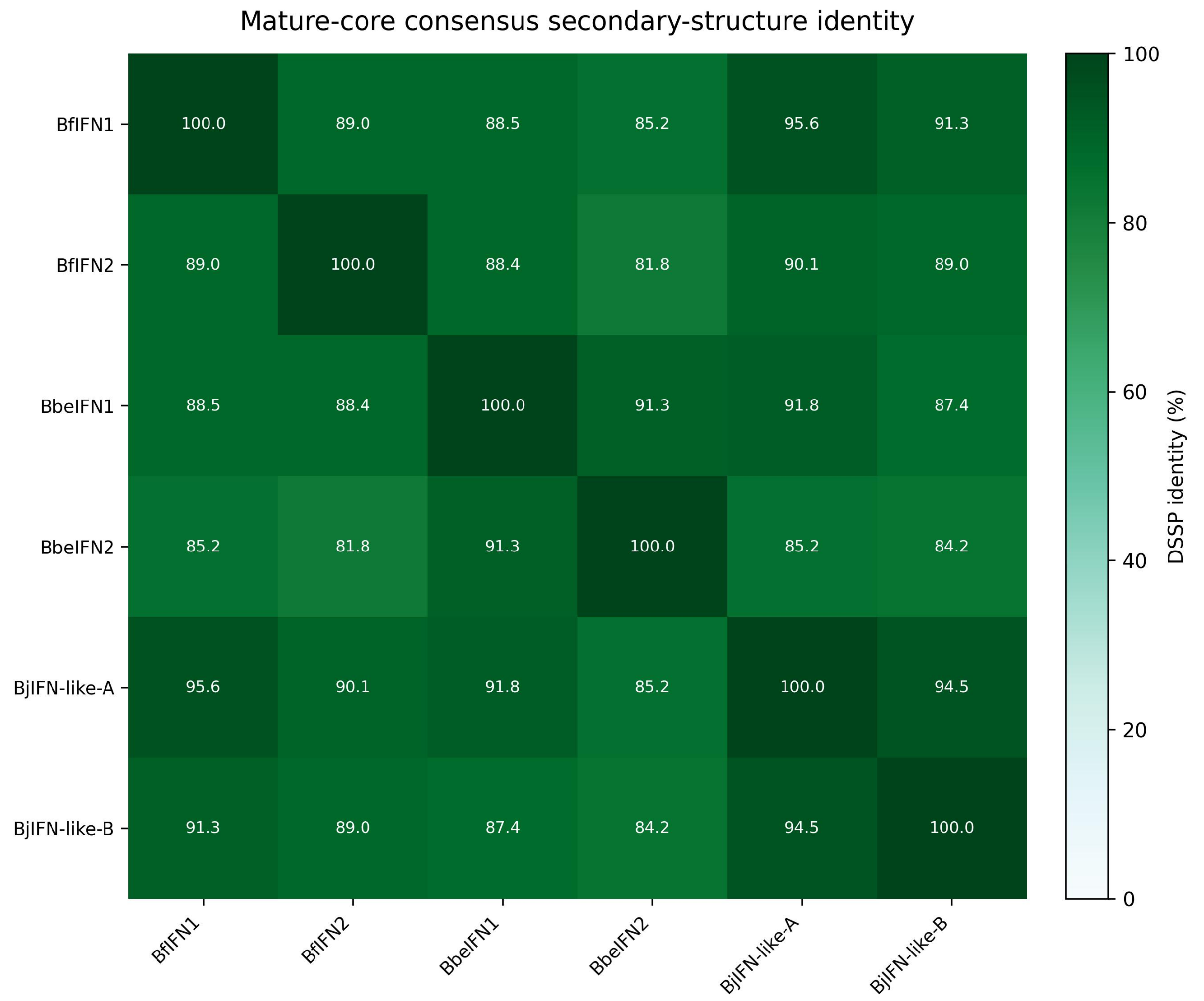


**Fig. S06: Mature-core consensus secondary structure identity of six amphioxus IFN-like proteins**. Heatmap illustrating pairwise DSSP-based secondary structure identity (%) calculated over the mature-core region of six amphioxus IFN-like proteins from *Branchiostoma floridae*, *Branchiostoma belcheri* and *Branchiostoma japonicum*. The colour bar denotes DSSP secondary structure identity ranging from 0% (white) to 100% (dark green). Numerical identity values are annotated within each cell.

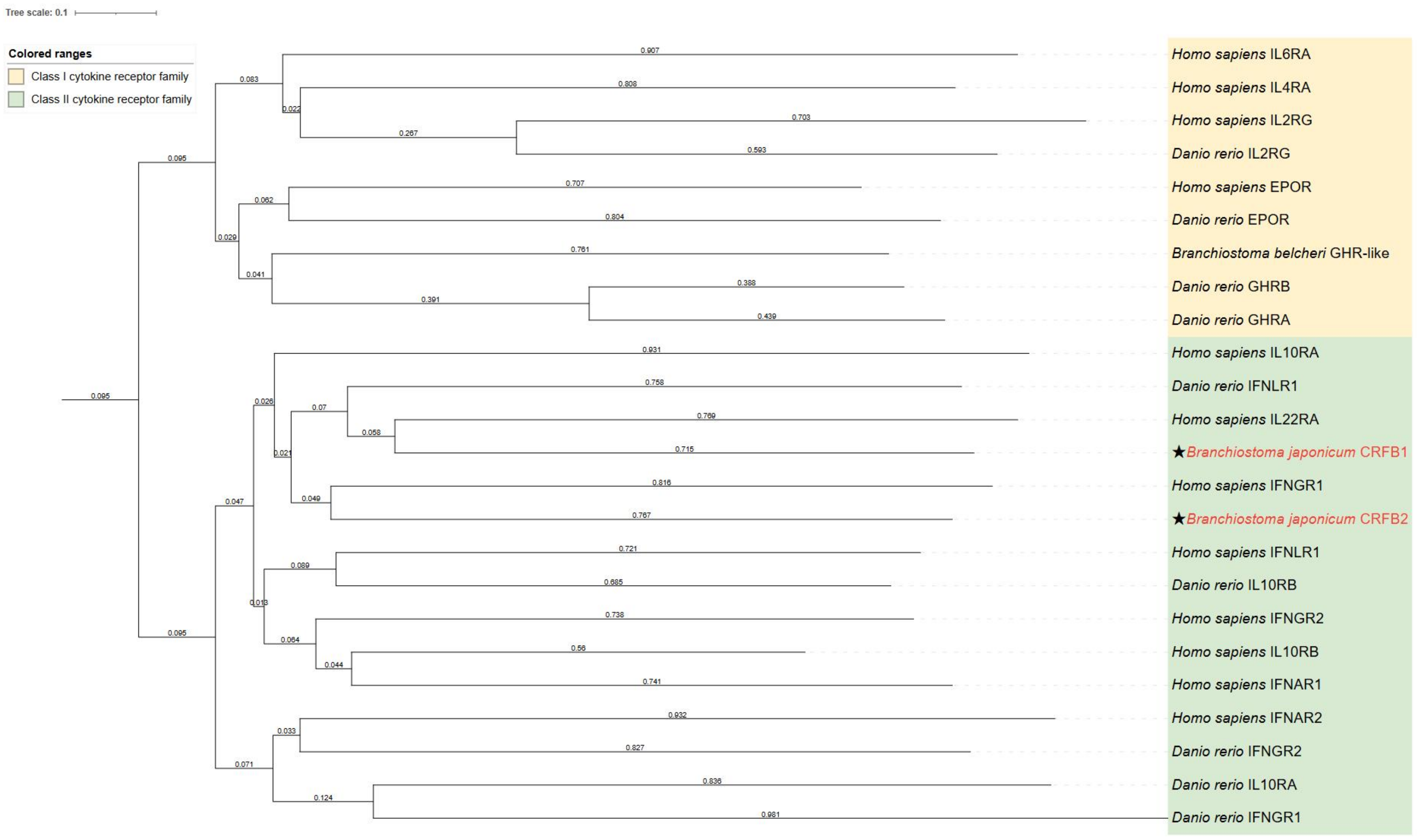


**Fig.S07: Phylogenetic analysis of amphioxus CRFB1&2 and class I and class II cytokine receptors in vertebrates**. The phylogenetic tree was constructed based on the amino acid sequences of the receptor extracellular domains (ECDs). The yellow-shaded region indicates the clade of the Class I cytokine receptor family, and the green-shaded region indicates the clade of the Class II cytokine receptor family. The black stars (★) mark the amphioxus receptors Branchiostoma japonicum CRFB1 and CRFB2 identified in this study, both of which are clustered within the Class II cytokine receptor family clade. The numbers on the branches represent the Bootstrap support values (1000 replicates).

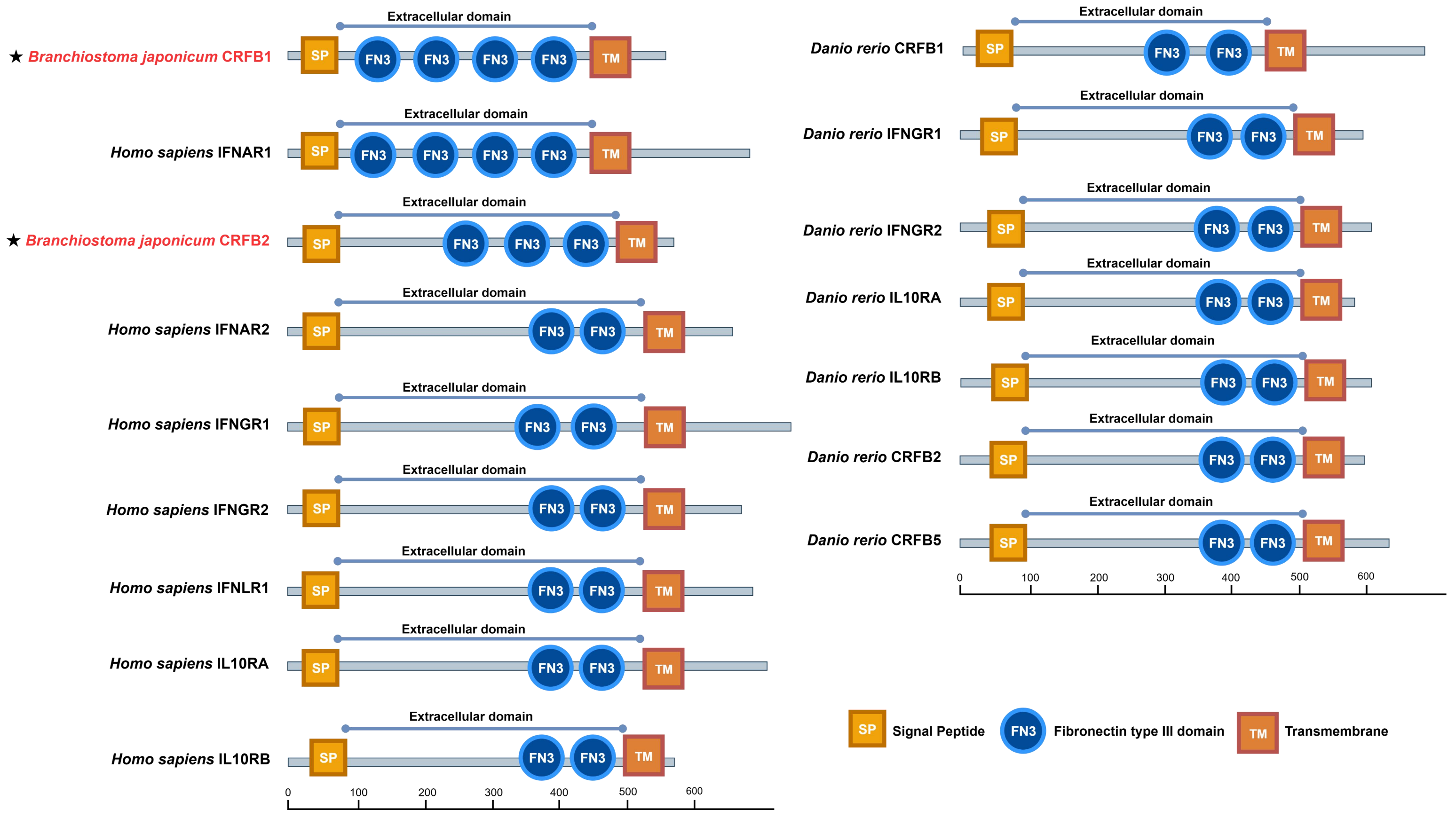


**Fig.S08: Comparison of domain architectures between amphioxus CRFB1/2 and vertebrate class II cytokine receptors.** Domain composition of extracellular regions of CRFB1/2 in Branchiostoma japonicum, Class II cytokine receptors in *Homo sapiens* and *Danio rerio.* SP (yellow box): Signal Peptide; FN3 (blue circle): Fibronectin type III domain; TM (red box): Transmembrane region. *B. japonicum* CRFB1 contains 4 FN3 domains, and CRFB2 contains 3 FN3 domains. Both exhibit the typical “signal peptide–FN3 domain module–single transmembrane region” topology of this receptor family.

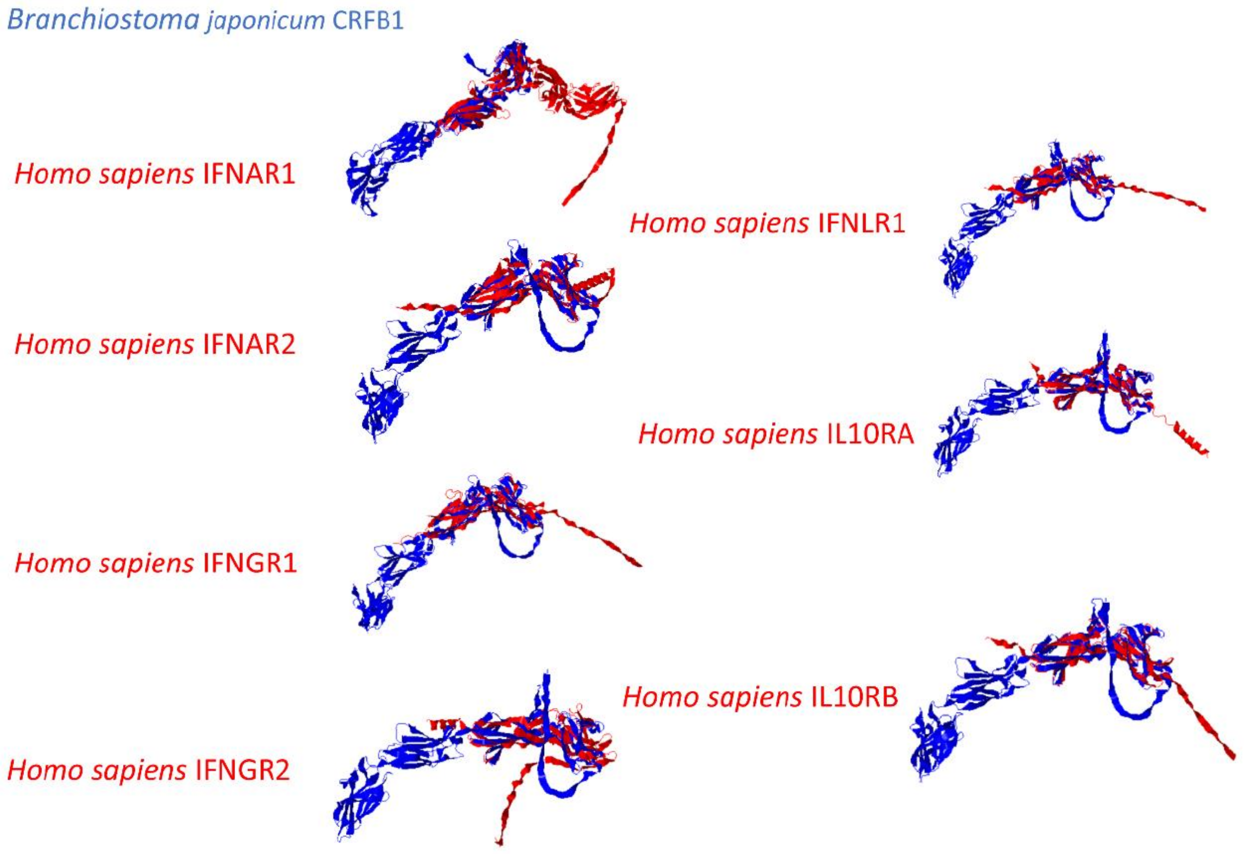


**Fig.S09: Three dimensional structural superimposition of the extracellular domain of amphioxus BjCRFB1 against human class II cytokine receptors.** Ribbon-view structural superimpositions of the extracellular domain (ECD) of *Branchiostoma japonicum* BjCRFB1 (blue) onto the extracellular domains of multiple human class II cytokine receptors (red), including IFNAR1, IFNAR2, IFNGR1, IFNGR2, IFNLR1, IL10RA and IL10RB. The conserved FN3 domain core architecture is observed, whereas flexible loop regions exhibit prominent structural divergence.

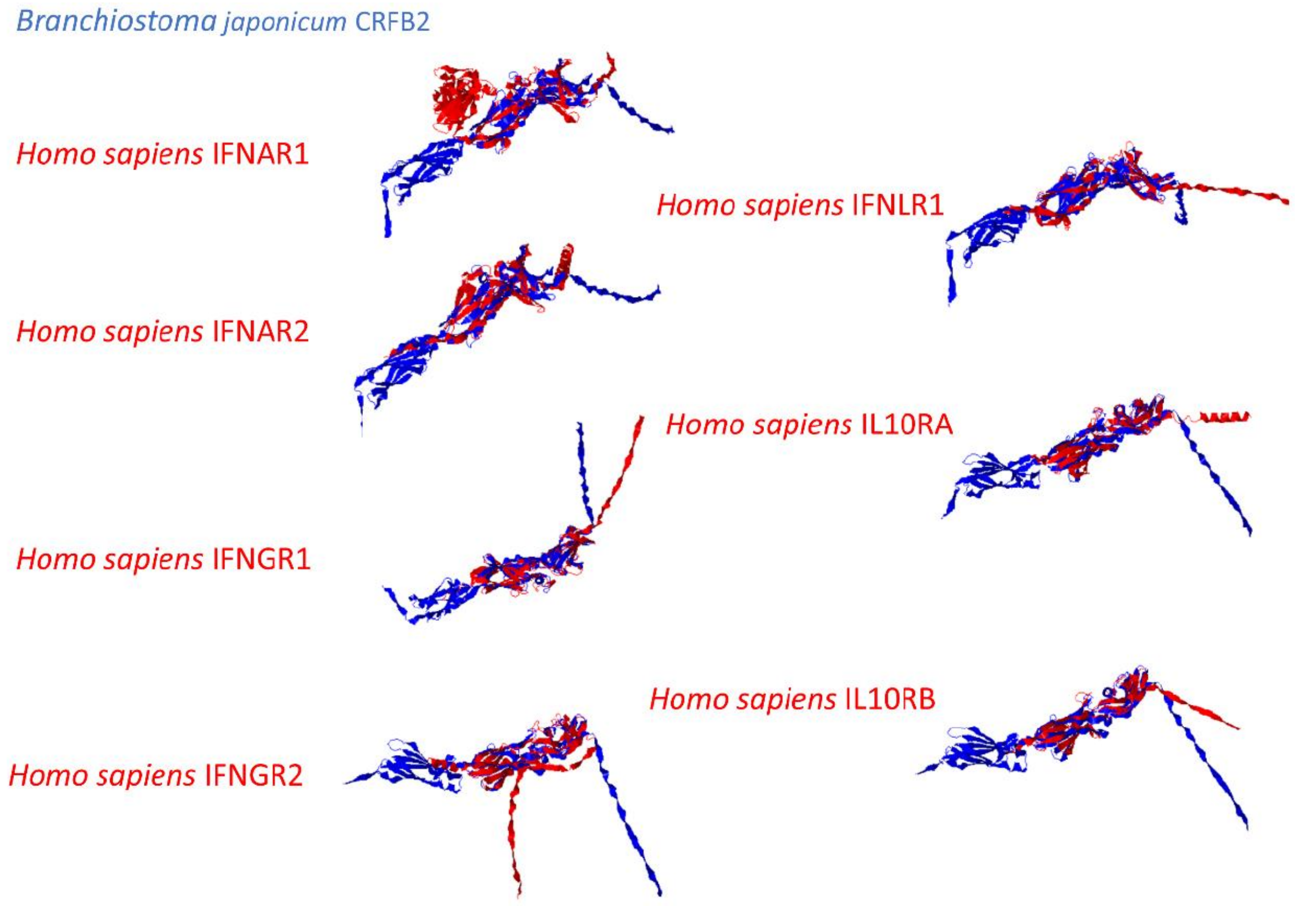


**Fig.S10: Three dimensional structural superimposition of the extracellular domain of amphioxus BjCRFB2 against human class II cytokine receptors.** Ribbon-view structural superimpositions of the extracellular domain (ECD) of *B. japonicum* BjCRFB2 (blue) onto the extracellular domains of human class II cytokine receptors (red), including IFNAR1, IFNAR2, IFNGR1, IFNGR2, IFNLR1, IL10RA and IL10RB. Conserved core folding of FN3 domains is observed, while flexible loop regions display obvious structural divergence. Compared with BjCRFB1 (Fig. S09), BjCRFB2 exhibits higher structural resemblance to human IFN-γ receptor subunits.

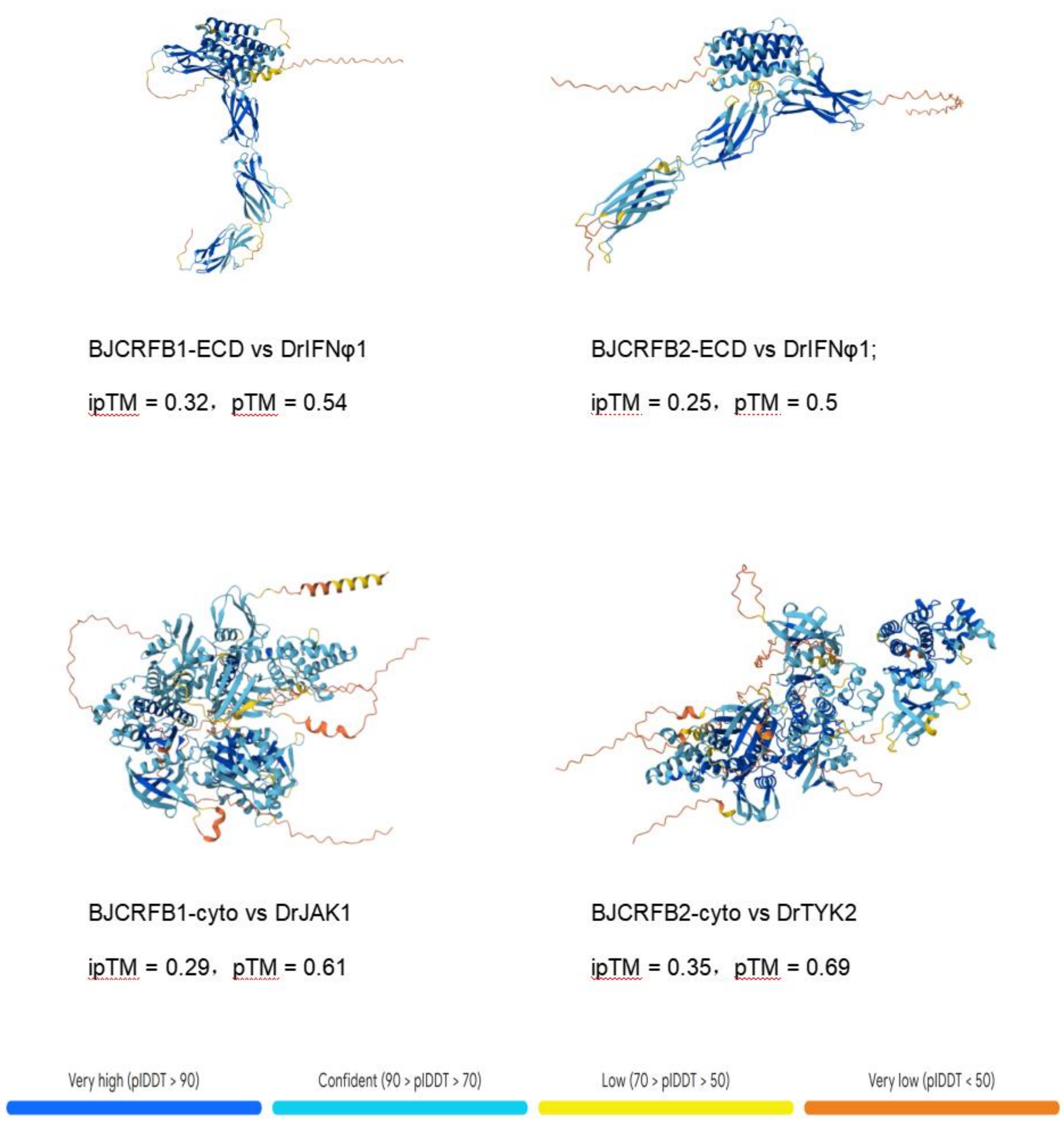


**Fig.S11: AlphaFold-based structural modelling of BjCRFB1 and BjCRFB2 with fish IFNφ and JAK–STAT-associated signalling proteins.** Predicted complex structures of BjCRFB1/BjCRFB2 with fish IFNφ and the JAK–STAT-related proteins DrJAK1 and DrTYK2 are shown. Structures are coloured according to the pLDDT confidence metric: very high (pLDDT > 90, dark blue), confident (70 < pLDDT ≤ 90, cyan), low (50 < pLDDT ≤ 70, yellow), and very low (pLDDT < 50, orange). The ipTM and pTM scores are indicated for each predicted complex. The relatively low ipTM scores indicate limited confidence in the predicted inter-chain interfaces; therefore, these models are interpreted as exploratory evidence of potential structural compatibility rather than direct evidence of physical interaction.

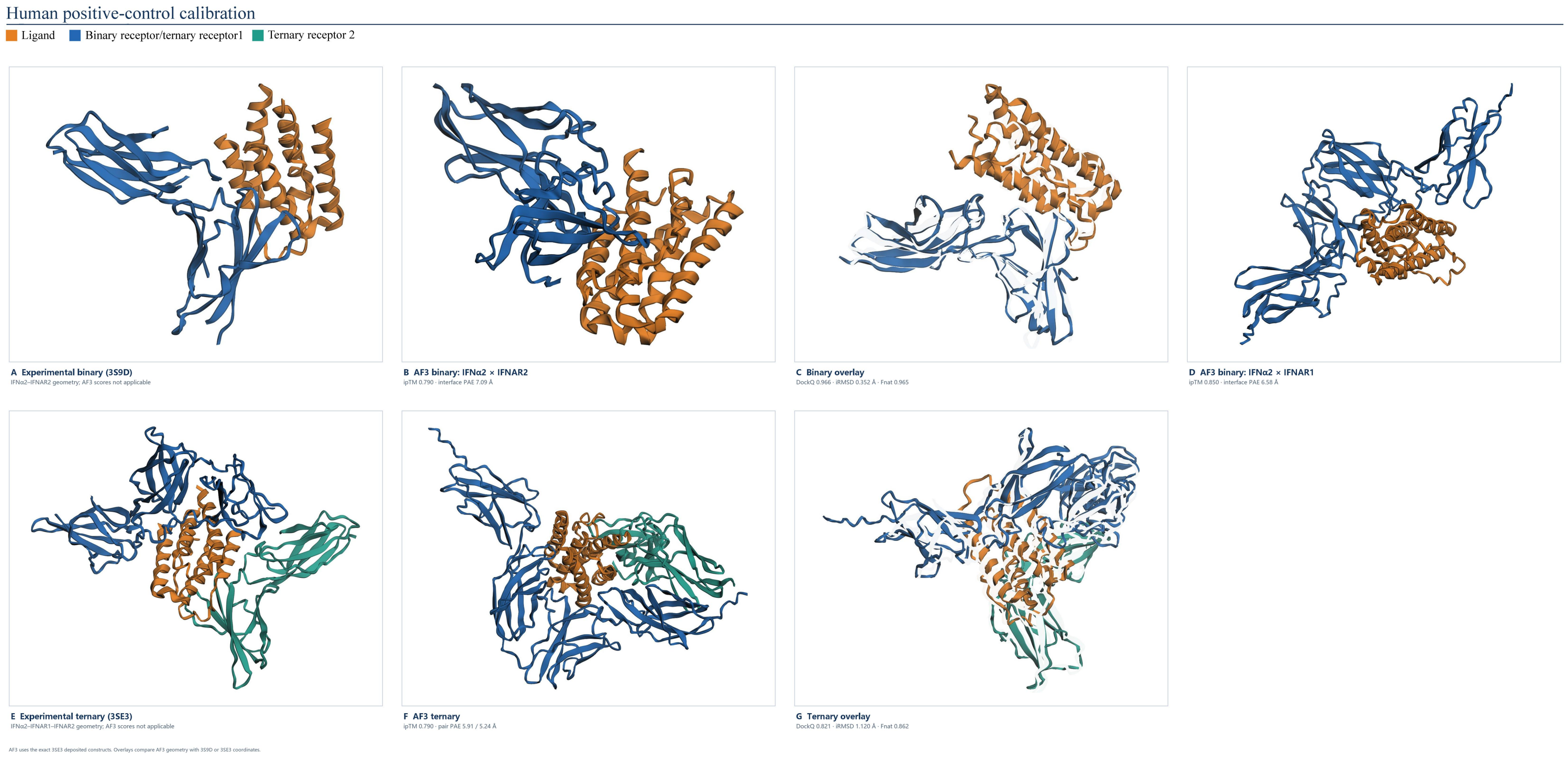


**Fig.S12: Human positive-control benchmark for protein-complex prediction.** AlphaFold3 complex prediction performance was benchmarked against experimentally determined human IFN-IFNAR binary (PDB: 3S5D) and ternary (PDB: 5T63) ligand-receptor complexes. **(A)** Experimentally solved binary IFNα2-IFNAR2 complex (PDB: 3S5D). **(B)** AlphaFold3 predicted binary IFNα2-IFNAR2 complex, with modelling metrics (ipTM and interface PAE) annotated below the panel. **(C)** Structural overlay superimposing the experimental binary structure (transparent white) onto the AlphaFold3 binary prediction, with DockQ, RMSD and Fnat values provided as quantitative geometric metrics. **(D)** AlphaFold3-predicted binary IFNα2-IFNAR1 complex. **(E)** Experimentally solved ternary IFNα2-IFNAR1-IFNAR2 complex (PDB: 5T63). **(F)** AlphaFold3-predicted ternary IFNα2-IFNAR1-IFNAR2 complex; corresponding ipTM, pair PAE and interface PAE values are indicated. **(G)** Structural overlay of the experimental ternary complex (transparent white) aligned to the AlphaFold3 ternary prediction, with DockQ, RMSD and Fnat reported for structural comparison. Colour coding: orange, IFNα2 ligand; blue, primary receptor subunit; green, secondary receptor subunit. This high-affinity vertebrate IFN-receptor complex serves as a reference baseline for evaluating amphioxus IFN-CRFB docking models.

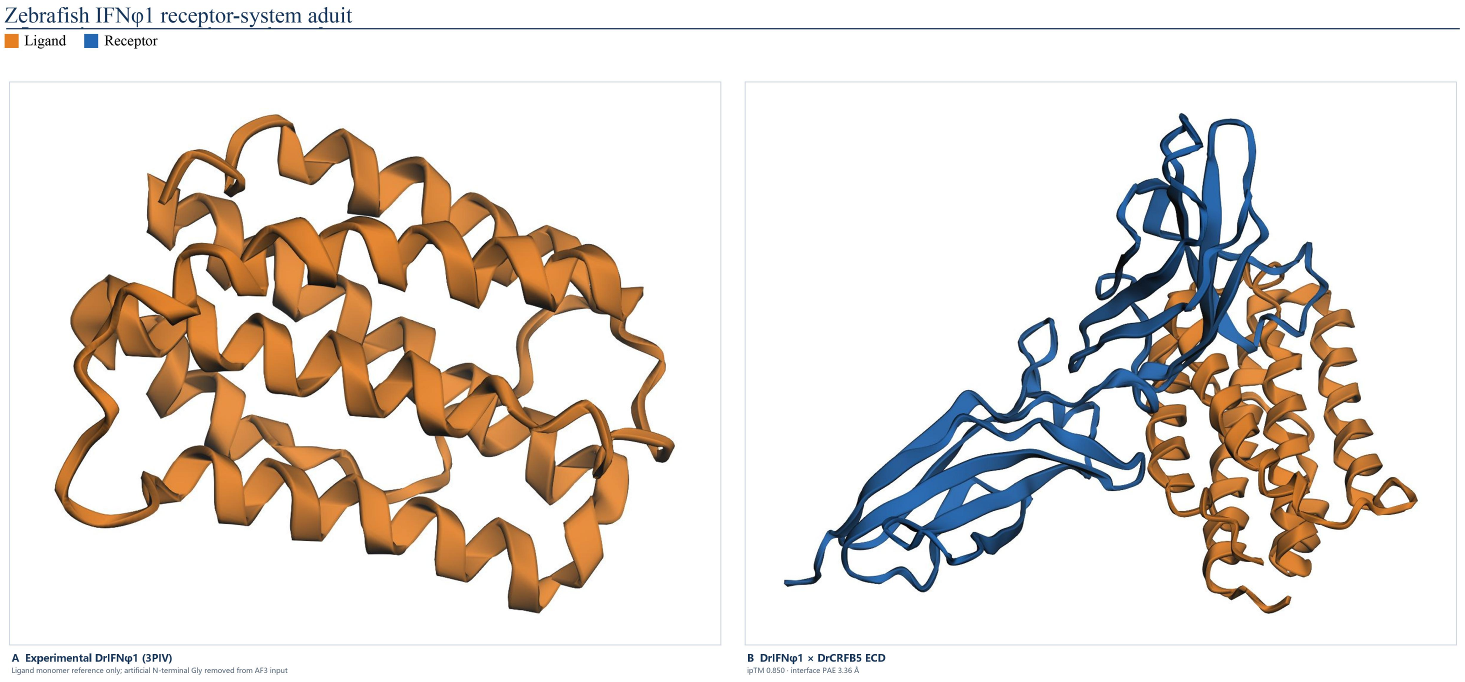


**Fig.S13: AlphaFold3 modelling of the zebrafish IFNφ1-CRFB5 receptor system. (A)** Crystal structure of DrIFNφ1 monomer (PDB: 3PIV). The artificial N-terminal glycine was trimmed before AlphaFold3 computation. Ligand is shown in orange. **(B)** AlphaFold3 predicted DrIFNφ1-DrCRFB5 extracellular domain complex. Orange represents DrIFNφ1 ligand, blue represents DrCRFB5 receptor ectodomain. Interface ipTM = 0.850, interface PAE = 3.36 Å.

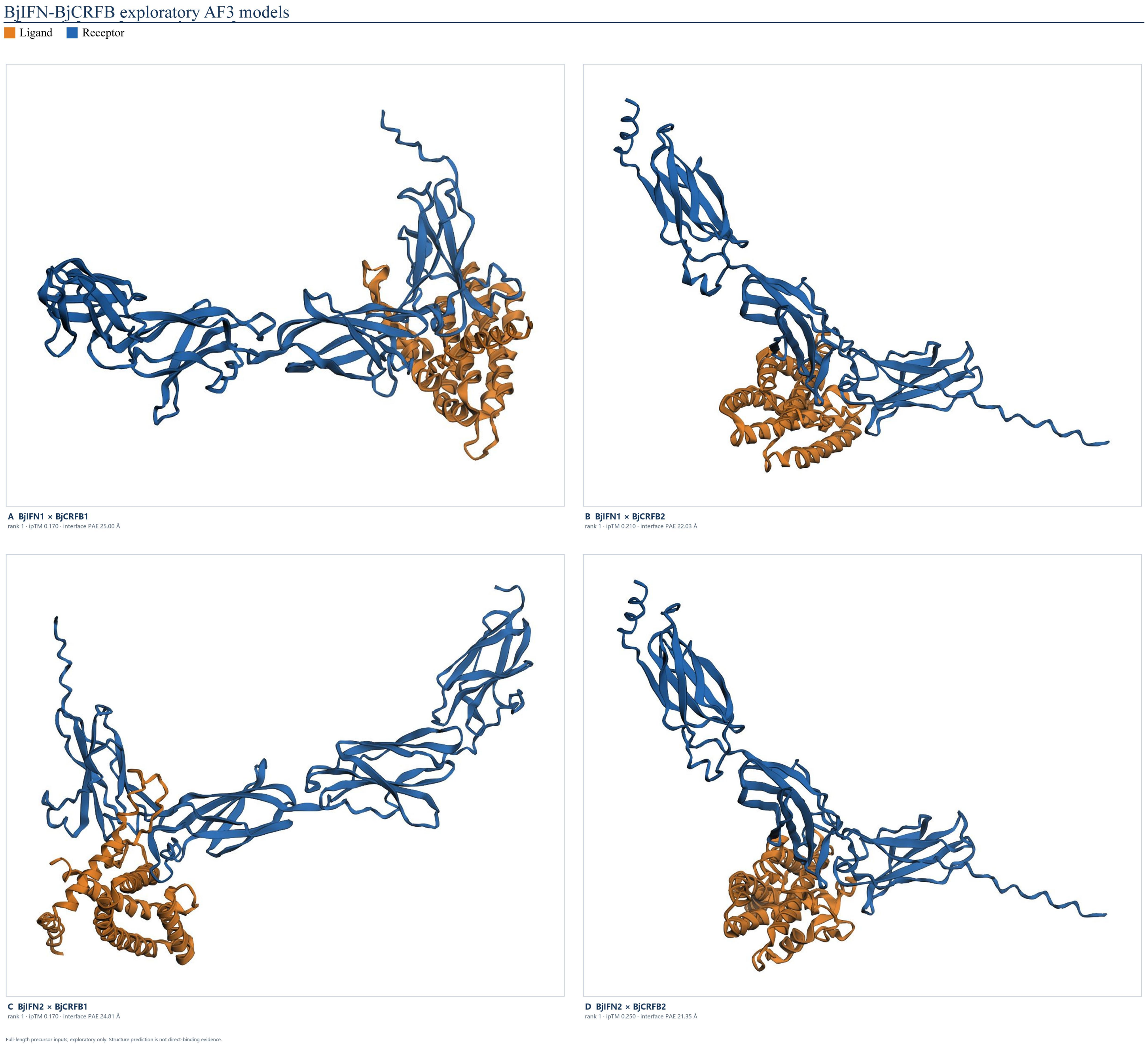


**Fig.S14. Four rank-1 full-length AlphaFold3 exploratory models of BjIFN-BjCRFB ligand-receptor complexes.** AlphaFold3 rank-1 full-length complex predictions for amphioxus IFN-CRFB pairs: **(A)** BjIFN1-BjCRFB1; **(B)** BjIFN1-BjCRFB2; **(C)** BjIFN2-BjCRFB1; **(D)** BjIFN2-BjCRFB2. Orange represents BjIFN ligands; blue represents BjCRFB receptors. Low ipTM and high PAE values across these models indicate primitive, low- confidence ligand- receptor interfaces. Among these pairwise combinations, the BjIFN2-BjCRFB2 complex shows relatively more favourable docking geometry, suggesting early stage ligand receptor binding preference specificity in amphioxus.

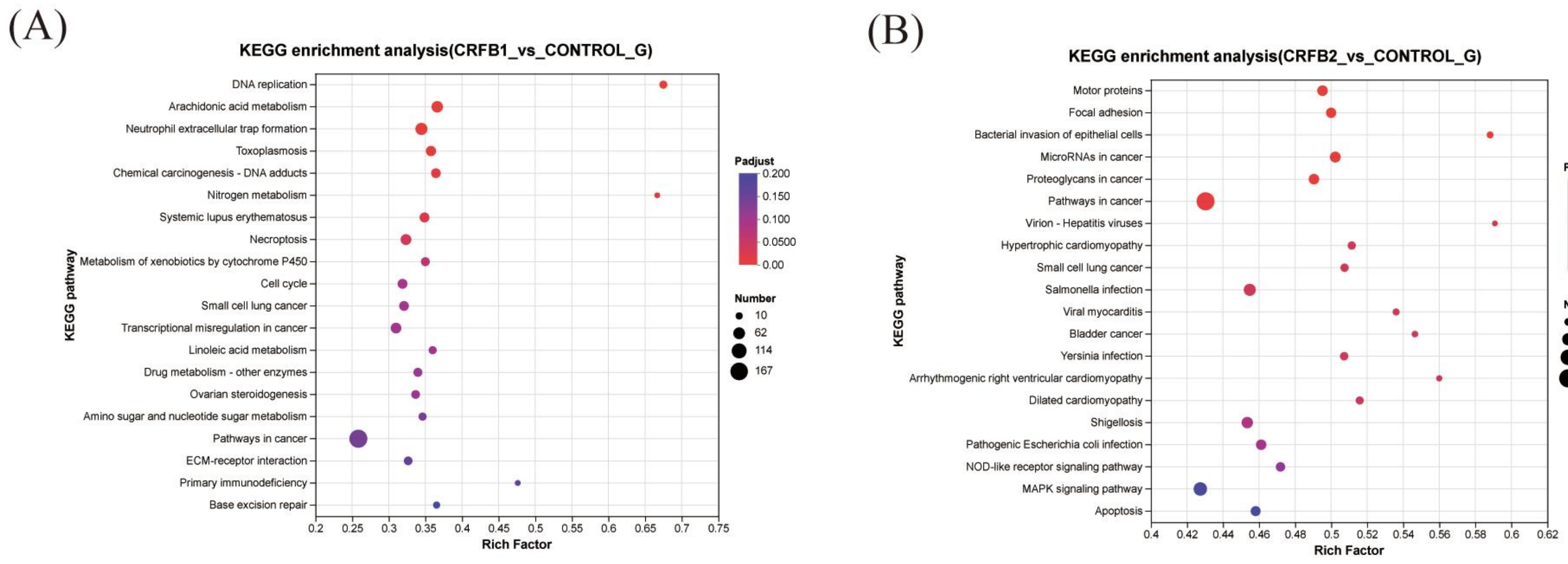


**Fig.S15**. **Transcriptome sequencing indicates BjCRFB1/2 regulate immune-related gene expression**. KEGG enrichment bubble plots for CRFB1-overexpression **(A)** and CRFB2-overexpression **(B)** transcriptome comparisons against control. X-axis: Rich factor; y-axis: KEGG pathway terms. Bubble colour: Padjust value; bubble size: count of differentially expressed genes. BjCRFB1 and BjCRFB2 drive distinct sets of enriched immune signalling related pathways, consistent with their immunomodulatory roles. Plot note: Lower Padjust values (red colour) represent higher statistical significance of enrichment.

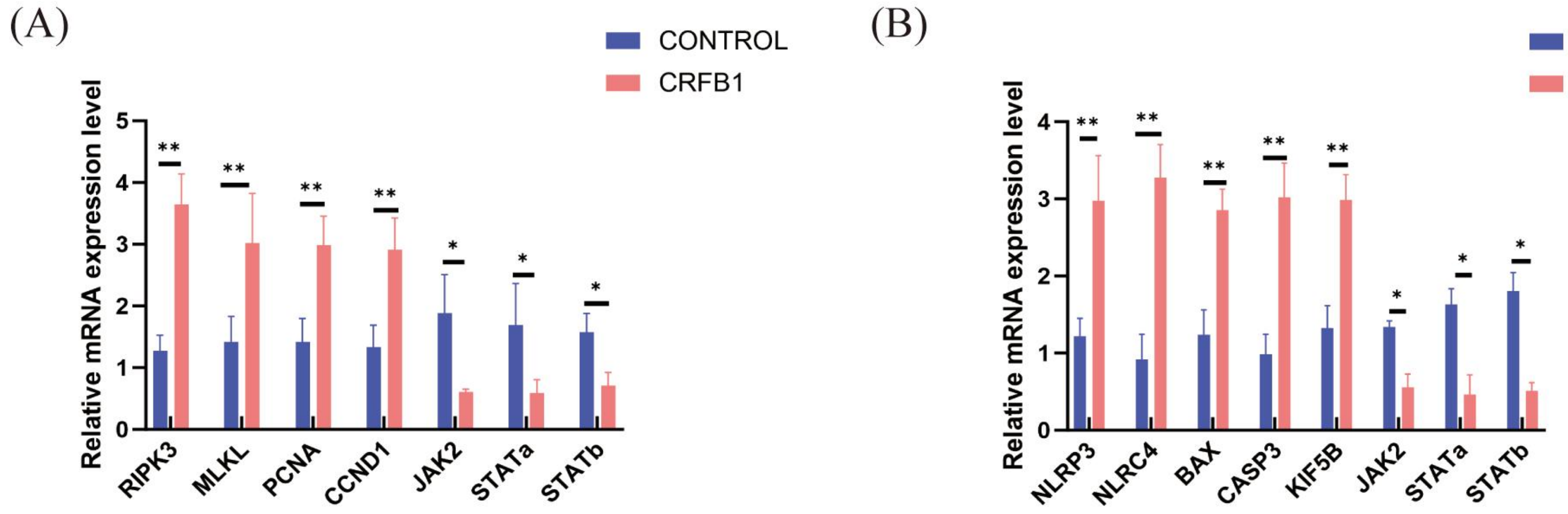


**Fig.S16. qPCR validation of immune-related-gene-expression modulated by BjCRFB1 and BjCRFB2.** qRT-PCR quantification of relative mRNA expression levels for selected target genes. **(A)** BjCRFB1 overexpression group versus control group. **(B)** BjCRFB2 overexpression group versus control group. Blue bars represent empty vector control group; red bars represent receptor overexpression group. Data are shown as mean ± SD. *$P < 0.05$, **$P < 0.01$. qPCR results independently confirm the transcriptome enrichment observations in Figure S15.

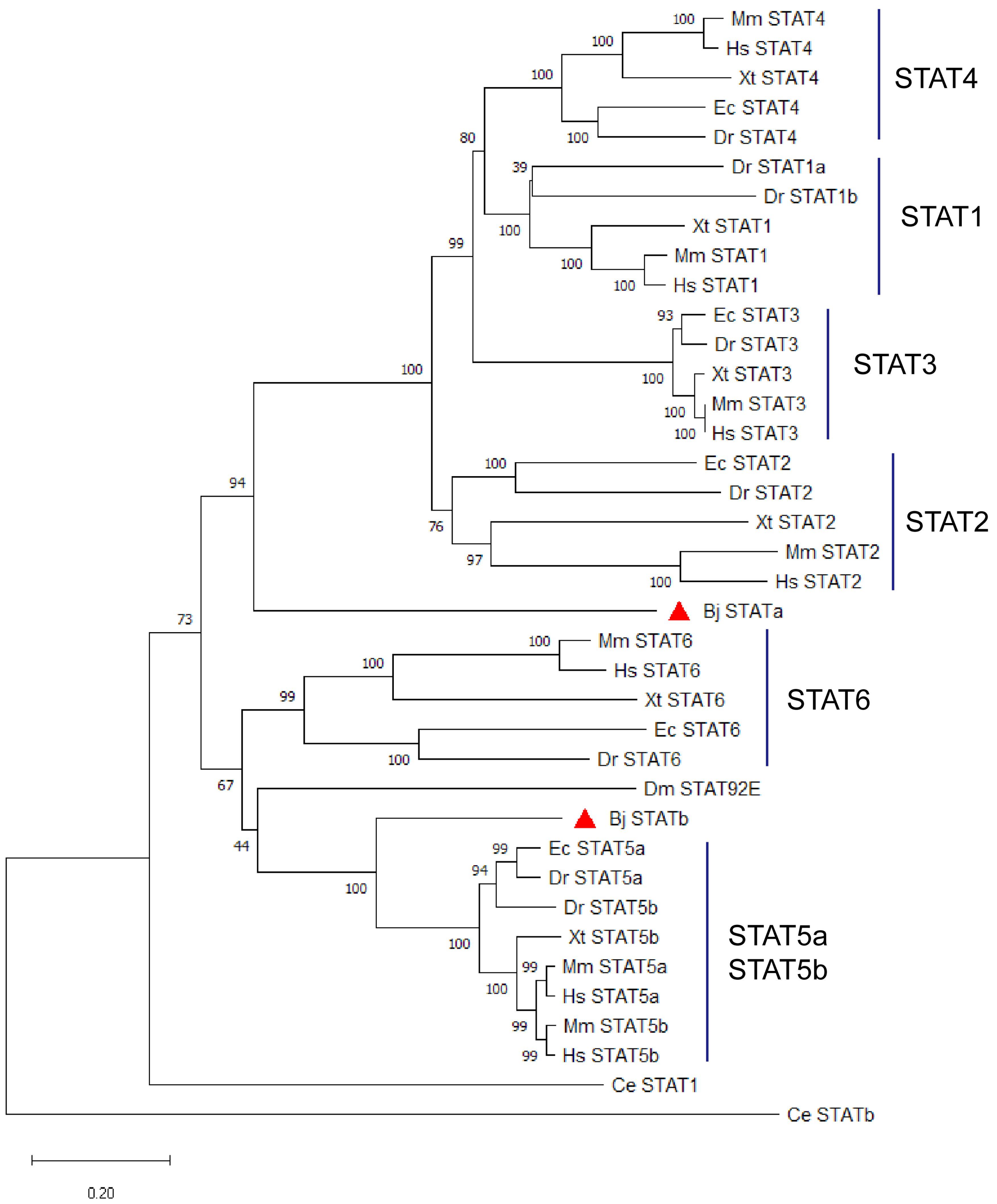


**Fig.S17: Maximum-likelihood phylogenetic tree of chordate STAT family proteins.** Phylogenetic reconstruction of STAT homologues across chordate lineages. BjSTATa clusters basal to the vertebrate STAT1-STAT4 subfamily (related to vertebrate STAT6); BjSTATb falls at the basal branch of vertebrate STAT5 clade. Bootstrap support values are labelled at internal nodes. Amphioxus STAT paralogs represent undifferentiated ancestral prototypes prior to vertebrate STAT-subtype diversification. Species abbreviation：Mm: *Mus musculus*；Hs: *Homo sapiens*；Xt: *Xenopus tropicalis*；Ec: *Erpetoichthys calabaricus*；Dr: *Danio rerio*；Bj: *Branchiostoma japonicum*；Dm: *Drosophila melanogaster*；Ce: *Caenorhabditis elegans*.

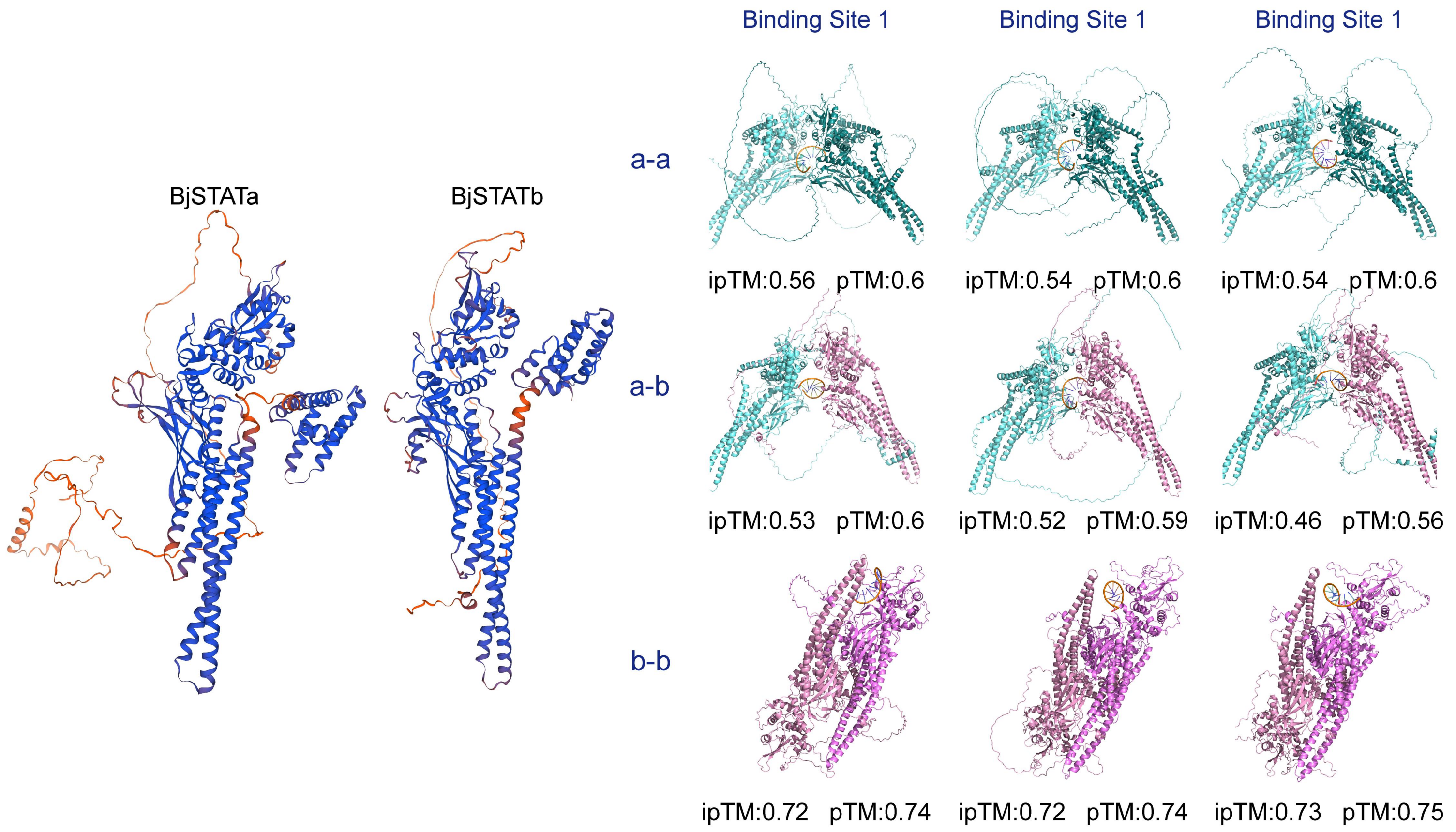


**Fig.S18: Structural docking predictions for BjSTAT homo- and heterodimers bound to viperin promoter cis-elements.** AlphaFold3 structural predictions of three BjSTAT dimer configurations: BjSTATa-BjSTATa homodimer (a-a), BjSTATa-BjSTATb heterodimer (a-b), and BjSTATb-BjSTATb homodimer (b-b), docked against three putative STAT-binding sites (Binding Site 1, 2, 3) within the *viperin* promoter. Left panels show monomer structures of BjSTATa and BjSTATb. Interaction metrics (ipTM and pTM scores) for each dimer-DNA complex are annotated beneath the corresponding structural models. The BjSTATb-BjSTATb homodimer exhibits the highest ipTM/pTM values, indicating superior predicted binding affinity, followed by the BjSTATa-BjSTATb heterodimer and the BjSTATa-BjSTATa homodimer. All three dimers are capable of engaging the predicted promoter-binding sites, supporting multiple combinatorial transcriptional regulatory modes downstream of BjCRFB-JAK-STAT signalling.

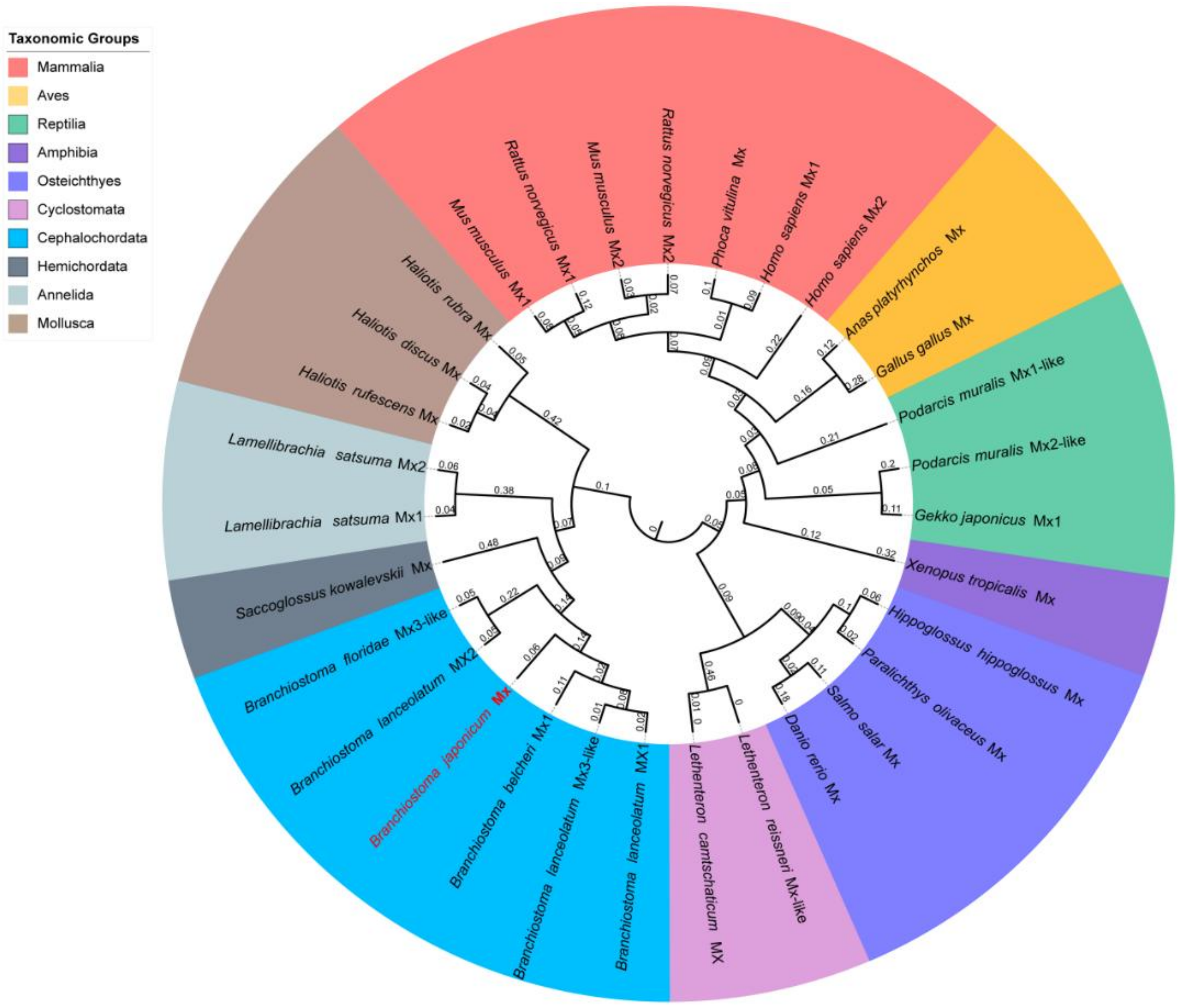


**Fig.S19: Phylogenetic analysis of Mx proteins from diverse metazoans**. Circular maximum-likelihood phylogenetic tree constructed using Mx amino acid sequences from representative metazoan species. Taxonomic clades are colour-coded as indicated. *Branchiostoma japonicum* BjMx is marked in red. Bootstrap values (1000 replicates) are shown on tree branches. This phylogeny places BjMx at the evolutionary boundary between invertebrate and vertebrate Mx homologs.

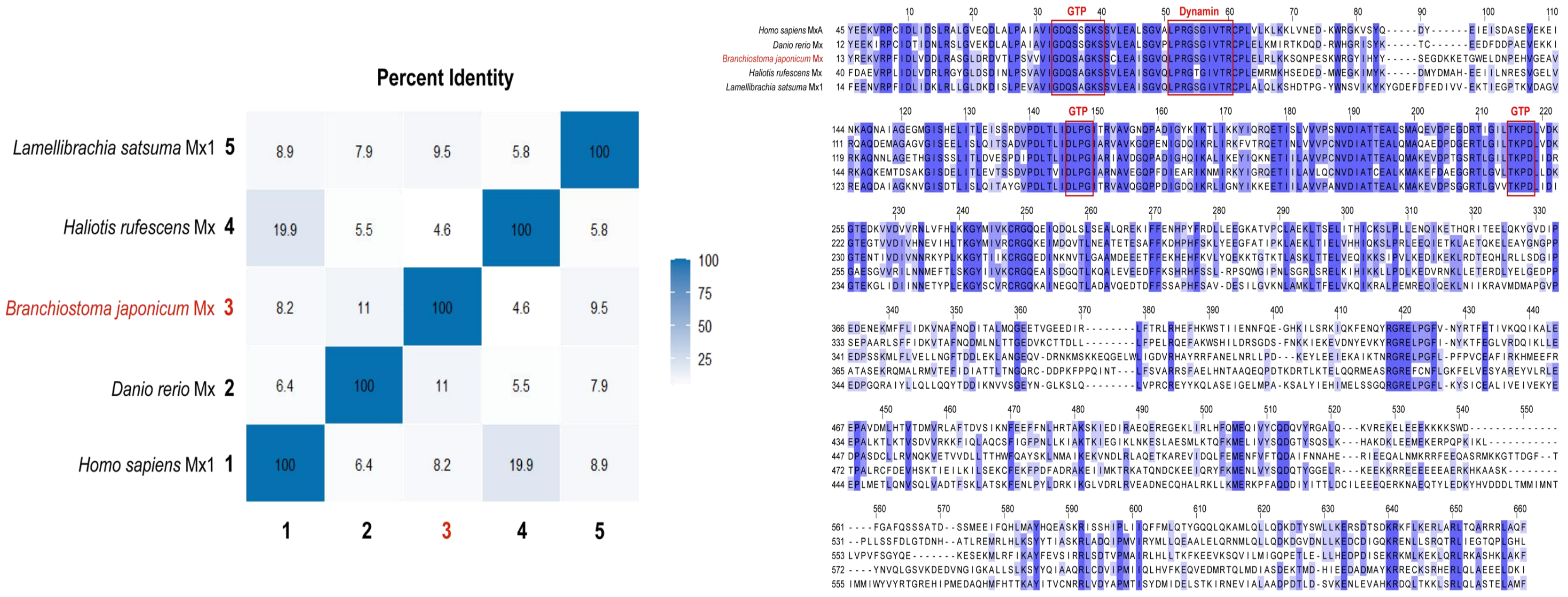


**Fig.S20: Sequence identity and multiple-sequence alignment of representative Mx proteins across metazoans**. Left: Heatmap showing pairwise percent-identity values among Mx protein sequences from *Homo sapiens*, *Danio rerio*, *Branchiostoma japonicum*, *Haliotis rufescens* and *Lamellibrachia satsuma*. Colour gradient indicates sequence identity from low (light-blue) to high (dark-blue). Right: Multiple sequence alignment of Mx homologs. Conserved GTP-binding and Dynamin-family functional motifs are boxed in red. Conserved amino acid residues are shaded in blue. Despite low overall sequence identity between BjMx and vertebrate Mx proteins, the core GTP-binding domains are highly conserved.

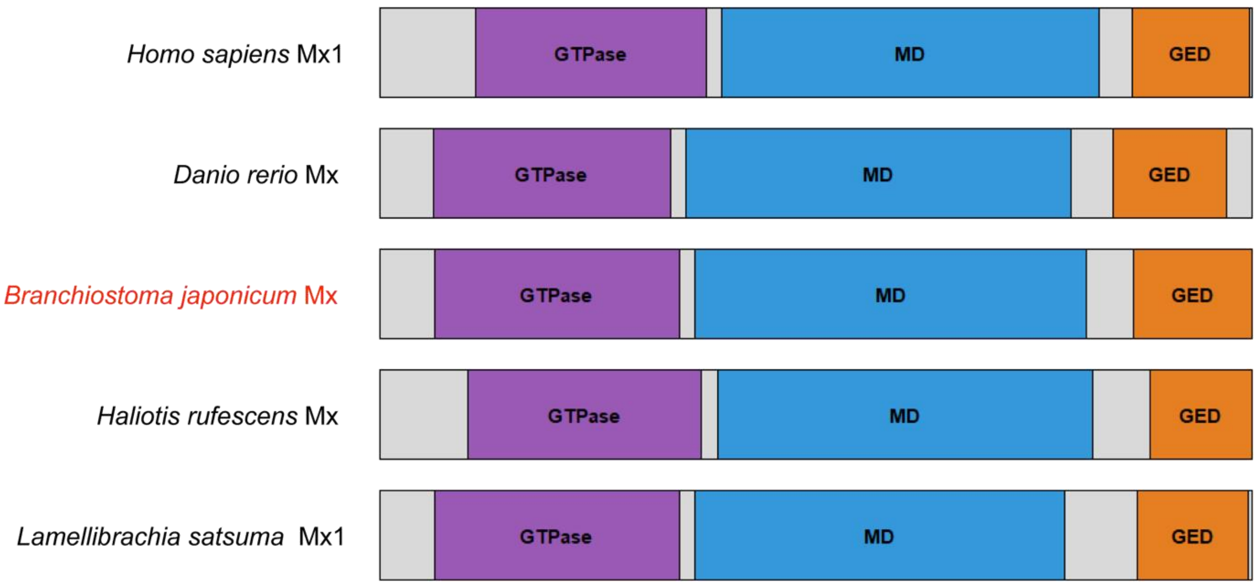


**Fig.S21: Domain architecture of Mx proteins from distinct metazoan species**. Schematic representation of domain organization of Mx homologs from *Homo sapiens*, *Danio rerio*, *Branchiostoma japonicum*, *Haliotis rufescens* and *Lamellibrachia satsuma*. Conserved modular domains are indicated: GTPase domain (purple), middle domain (MD, blue), and GTPase-effector domain (GED, orange). BjMx is highlighted in red. The domain composition reveals that BjMx harbours the canonical tripartite domain architecture characteristic of vertebrate Mx GTPases.

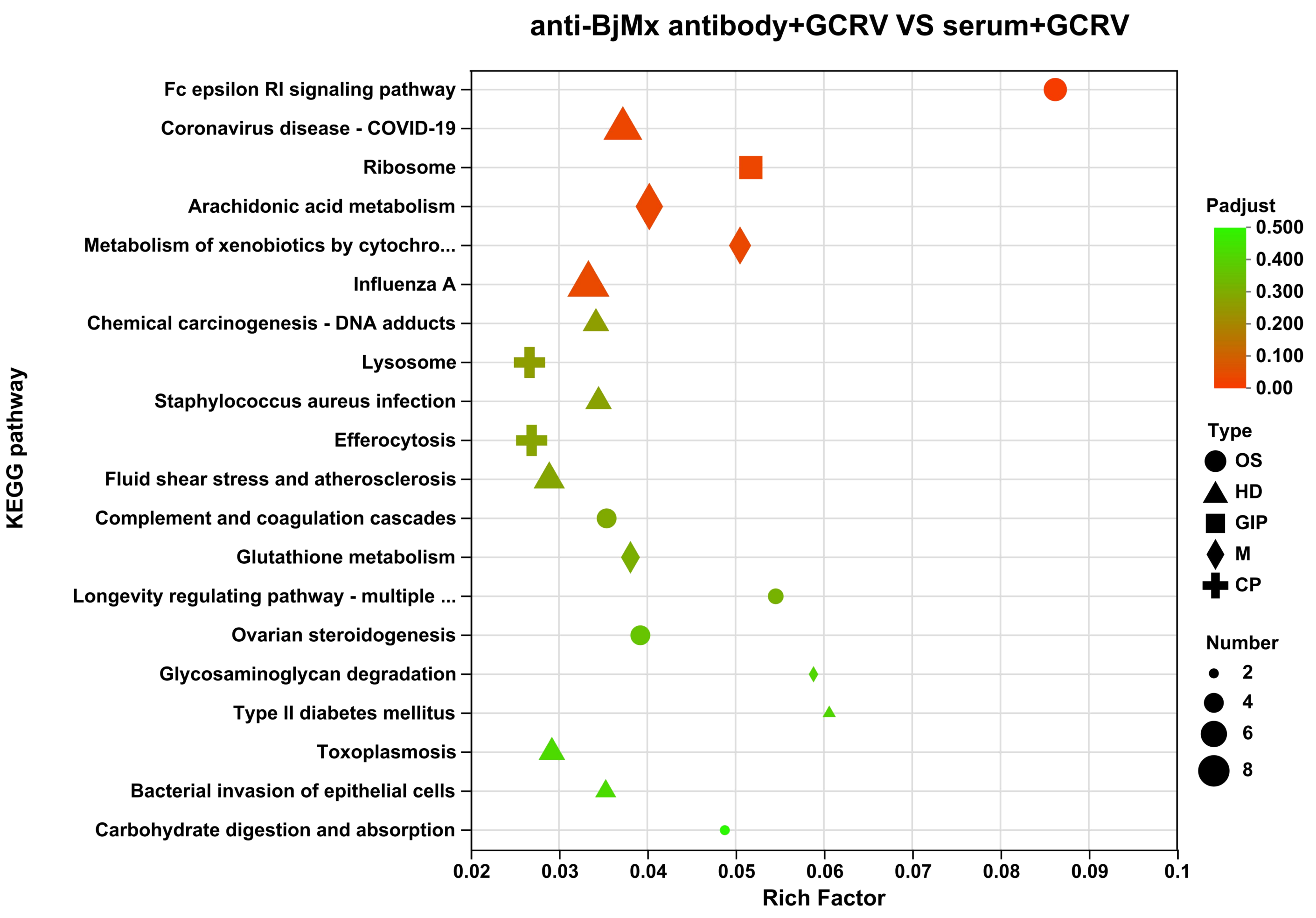


**Fig.S22: KEGG pathway enrichment analysis for BjMx loss of function group upon GCRV challenge**. Bubble plot of KEGG pathway enrichment derived from transcriptomic comparison between the anti-BjMx antibody treated group and serum control group following GCRV challenge. X-axis represents the Rich factor; colour gradient corresponds to adjusted *P*-value (Padjust); symbol size indicates the number of enriched differentially expressed genes. Antibody-mediated neutralization of endogenous BjMx triggers compensatory broad-spectrum antiviral responses, over-activated innate immune signalling (e.g. Fc epsilon RI signalling pathway), accompanied by dysregulated arachidonic acid mediated inflammatory lipid metabolism and xenobiotic detoxification responses (cytochrome P450-related pathways).
